# Explainable Transformer Models for Functional Genomics in Prokaryotes

**DOI:** 10.1101/2020.03.16.993501

**Authors:** Jim Clauwaert, Gerben Menschaert, Willem Waegeman

## Abstract

The effectiveness of deep learning methods can be largely attributed to the automated extraction of relevant features from raw data. In the field of functional genomics, this generally comprises the automatic selection of relevant nucleotide motifs from DNA sequences. To benefit from automated learning methods, new strategies are required that unveil the decision-making process of trained models. In this paper, we present several methods that can be used to gather insights on biological processes that drive any genome annotation task. This work builds upon a transformer-based neural network framework designed for prokaryotic genome annotation purposes. We find that the majority of sub-units (attention heads) of the model are specialized towards identifying DNA binding sites. Working with a neural network trained to detect transcription start sites in *E. coli*, we successfully characterize both locations and consensus sequences of transcription factor binding sites, including both well-known and potentially novel elements involved in the initiation of the transcription process.

## 1 Introduction

Deep learning techniques are increasingly obtaining state-of-the-art performances for a multitude of prediction tasks in genomics (1). Capable of automating the feature extraction process during training, these techniques flip the conventional approach of machine learning studies; instead of requiring meaningful descriptors that function as input to the model, the raw input data is used. As a result, the decision-making process of the trained models needs to be examined in order to extract biological interpretation from them.

Deep learning is increasingly seen as a valid candidate to offer solutions were the current understanding of biological processes is insufficient. As such, explainable models that can offer insights into the biological process for which the model is trained are likely to play a role in the discovery of new regulatory mechanisms.

Today, the main contributions towards dissecting the workings of neural networks are achieved by mapping the importance of input features towards the model output. However, existing techniques are limited as they only evaluate the influence of the input sequence on the model output. It is likely that such techniques are unable to reflect complex mechanisms between multiple regulatory elements involved in many biological processes.

In a previous study, we introduced a model for performing annotation tasks on the genomic sequences of prokaryotes, achieving state-of-the-art results for the identification of transcription start sites (TSSs), translation initiation sites and methylation sites. This model is based on the transformer-XL architecture, first introduced for natural language processing (2). Vaswani et al. (3) found that the attention heads, specialized units making up the core mechanism of the neural network, map relations that convey semantic meaning. Based on this finding, we aim to introduce new ways of extracting biological meaning from a neural network trained on annotating the genome.

This work offers several contributions. (i) We evaluate and compare the annotations of a variety of recent, mostly sequencing based, *in vivo* methodologies for the detection of transcription start sites applied on *E. coli*. Based on four independent data sets, we curate an improved set of annotations. (ii) We lay out our approach to characterize the function of attention heads. Many of these are specialized towards detecting the presence of regulatory elements based on sequence and positional information. (iii) Using this information, we offer a comparison with existing knowledge on the mechanisms involved during the transcription process. As such, we link existing findings to literature and discuss their relevance. (iv) Using the model output predictions along the full genome sequence, we get a better understanding into the model characteristics and potential flaws. Based on these results, we conclude that the analysis of trained transformer-based models for genome annotation tasks offer singular potential for extracting biological meaning, where the discussed techniques are applicable for any genome annotation task.

## 2 Related Work

Research and development of tools to facilitate the interpretation of nucleotide sequences goes back as far as 1983, with Harr et al. (4) describing mathematical formulas on the creation of a consensus sequence. Other strategies have been designed in order to create weight matrices (5) and rank alignments (6; 7). The increased knowledge in the field of molecular biology led towards feature engineering efforts of the nucleotide sequence. Such descriptors include, but are not limited to: the GC-content of sequences, bendability (8), flexibility (9) and free energy (10). Recently, Nikam et al. (11) published Seq2Feature, a tool that can extract 41 DNA sequence-based descriptors.

Despite efforts invested into linking patterns between engineered sequence features and genomic sites of interest, this objective has not been solved adequately, even with the help of early machine learning methods. Today, the branch of deep learning is increasingly returning state-of-the-art performances for a variety of tasks in the field of functional genomics (12). By automating the feature extraction process during training, it is possible for models to map previously undiscovered relations within the raw DNA sequence. However, in contrast to earlier applications, to understand the relation of biologically-relevant descriptors and biological processes, a look into the decision-making process of the trained network is required.

The most popular group of strategies seeks to map the sensitivity of the input towards the output. The first way of measuring this is by permutations of the input features (13). More recent permutation strategies are the work of Fisher et al. (14) and Zintgraf et al. (15). Another approach seeks to investigate the partial derivatives of the model output class with respect to the input features. The technique was first discussed by Simonyan et al. (16) in 2014. Further developments in this branch include the work of Sundararajan et al. (17) and Shrikumar et al. (DeepLIFT) (18).

In the combined fields of genomics and deep learning, ideas borrowed from studies on sensitivity analysis have been applied occasionally. For example, Alipanahi et al. (19) perform a sensitivity analysis to guide the construction of motifs with which the contribution of positional nucleotides towards the presence of protein binding sites is visualized on the DNA. The sensitivity of nucleotides has been evaluated towards the methylation state by Angermueller et al. (20). Hill et al. (21) use permutation of the input sequence to investigate the influence on translation initiation and termination sites. By applying a pairwise mutation map, the correlation of two positional features is shown. DeepLIFT was used for the construction of motifs around splice sites in Eukaryotes (22).

## 3 MATERIAL AND METHODS

### 3.1 Data sources

To reduce the influence of potential factors of (systematic) noise, the annotations of a variety of high-precision *in vivo* experimental methods were compared. In recent years, several experimental procedures have been developed in quick succession for the detection and annotation of TSSs in prokaryotes: Cappable-seq by Etwiller et al. (24), retrieving 16,348 TSSs, Single Molecule Real Time Cappable-seq (SMRT-Cappable-seq), by Yan et al., (25), retrieving 2,311 TSSs, and Simultanuous 5’ and 3’–end sequencing (SEnd-seq), by Ju et al. (26), retrieving 4,026 TSSs.

Given its importance to the community, the TSS annotations featured by RegulonDB are also included. RegulonDB features an up-to-date collection of manually curated and automated (dRNAseq) TSSs from a plethora of independent sources (23). It has been the chosen data set for several recent machine learning tasks aimed at identifying TSSs (27; 28; 29; 30). The positive set evaluated are the positions marked with ‘strong evidence’, summing to a total of 6,487 annotated positions. An overview of the data is listed in Table 1.

**Table 1:** Overview of the data sets properties used in this study. From left to right: data sets source, *in vivo* next generation sequencing technique, number of positive and negative labels and the partition of TSS shared with at least one other data sets. The annotations are derived or mapped (Cappableseq) on *E. coli* MG1655 (accession: *NC_000913.3’*). The total genome consists of 9,283,304 nucleotides.

| data sets source | Technique | Year | Positive labels | Negative labels | Shared TSSs |
| --- | --- | --- | --- | --- | --- |
| RegulonDB (23) | Collection | - | 6,487 (0.07%) | 9,276,817 (99.93%) | 2,476 (38.17%) |
| Etwiller et al. (24) | Cappable-seq | 2016 | 16,348 (0.18%) | 9,266,956 (99.82%) | 4,631 (28.33%) |
| Yan et al. (25) | SMRT-Cappable-seq | 2018 | 2,574 (0.03%) | 9,280,730 (99.97%) | 2,311 (89.78%) |
| Ju et al. (26) | SEnd-seq | 2019 | 5,502 (0.06%) | 9,277,802 (99.94%) | 4,026 (73.17%) |

### 3.2 Model architecture

A transformer-based neural network is used. This architecture is borrowed from our previous work, featuring a performance-based comparison with existing deep learning methods (31). This model evaluates the full genome sequence by processing it in consecutive segments of length *l*. Every input nucleotide *x* ∈ {*A, C, G, T*} is first transformed into a vector embedding ***h***^(0)^, after which it is transformed *k* times through addition (residual connection) with another vector, obtained by the multi-head attention function present in each layer. The final layer sends the output of the last hidden state ***h***^(*k*)^ through a fully-connected layer in order to obtain the model output. For each residual block, the vector that is summed with the input (to obtain ***h***^(1)^,…, ***h***^(*k*)^) is calculated using the hidden states of *l* upstream positions. The connections displayed in Figure 1 illustrate the inputs used to calculate each intermediary value within the network. If required, hidden states from the previous segment (*s* − 1) are accessible for the calculation of the new hidden states in segment *s*.

**Figure 1:**
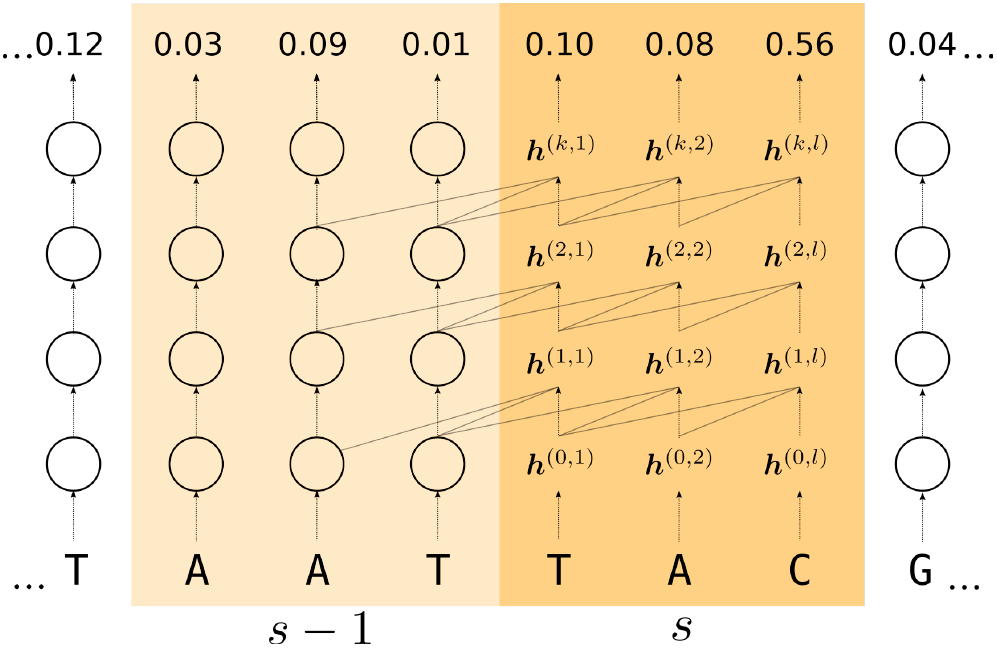
Illustration of the connectivity of intermediary values of the transformer architecture. The full genome is processed in sequential segments *s* with length *l*. First, the input nucleotide is transformed into a vector embedding ***h***^(0)^, after which it is processed by *k* consecutive residual blocks (***h***^(0)^ →… → ***h***^(*k*)^). The output probability is obtained by sending the final hidden state ***h***^(*k*)^ through a set of fully-connected layers. For the calculation at each residual block, the last *l* hidden states of the previous layer are applied. For example, ***h***^(1,*l*)^ is calculated by using the hidden states [***h***^(0,1)^,…, ***h***^(0,*l*)^]. Hidden states from the previous segment (*s* − 1) are made accessible for the calculation of the hidden states in segment *s*.

The multi-head attention applied in each residual block is methodologically identical. From each input hidden state, a query (***q***), key (***k***) and value (***v***) vector of equal shapes are calculated. The output ***z*** of the attention head applied on the hidden state at position *n* on the genome is calculated as follows:

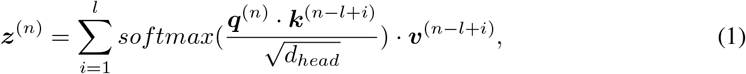

where the ***k*** and ***v*** vectors derived from *l* upstream hidden states are used. The multiplication of the query and key vector results in a score that is normalized for all input values using the *softmax* function. The denominator is used to stabilize the scores based on the dimensions of ***q, k*** and ***v*** (*d_head_*). The multiplication and normalization of the ***q*** and ***k*** vectors result in a set of attention scores that is attributed to the ***v*** vectors for the calculation of ***z*** (i.e. a linear combination). Essentially, the set of scores isolates the vectors of relevancy for that attention head, and is related to the information passed by ***v***. In each residual block, multiple attention heads are present (hence multihead attention), each featuring their own unique sets of scores to calculate ***q***, ***k*** and ***v***. As such, multiple types of information can be extracted from the input hidden states. The attention scores of the different attention heads are further processed into a single vector, which is summed with ***h*** to obtain the hidden state of the next layer (e.g. ***h***^(1)^ → ***h***^(2)^). A detailed overview of the model architecture and the mathematical operations performed are given in the Supplementary File.

In a previous study (31), we found that the contextual information embedded within the hidden states derived from single nucleotides is limited. As motifs are generally of greater importance towards biological processes, the addition of a convolutional layer results in the ***q, k*** and ***v*** vectors to be derived from multiple neighboring hidden states without affecting the input/output resolution. Thereby, the retrieval of relevant information using attention is improved, resulting in improved predictive performances on a variety of tasks (31).

Positional information is required within the vectors ***q, k*** and ***v***, as these are not implicit through the architecture of the network. This is achieved by superimposing (i.e. through summation) a positional encoding vector to ***h***. The added signal is in function of the vector index and the relative positioning with respect to the other input hidden states. It was shown that the transformer model is able to extract this information well and use it to obtain better performances (3; 2).

A model architecture was optimized for the detection of transcription start sites for a previous study, resulting in a model with a segment length of 512 nucleotides, 6 layers (i.e. residual blocks) and 6 attention heads within each residual block (31). To include information downstream of the annotated transcription start site towards its annotation, labels can be shifted downstream during processing times in order to include this information (i.e. position upstream) for each hidden state within the segment. In accordance to recent publications on the detection of TSSs, the downstream bound is positioned 20 nucleotides downstream of the transcription start site.

### 3.3 Training and evaluation

The transformer-based model was trained using the full genome sequence, resulting in a total sample size of 9,283,204. The sizes of the positive and negative samples are listed in Table 1. As the model iterates the genome sequentially, the training, test and validation set are created by splitting the genome at three positions that constitute 70%, 20% and 10% of the genome, respectively. The training, test and validation sets have equal input and output class distributions. In order to compare all data sets on the same genome, annotations of the Cappable-seq experiment, originally mapped on the *U00096.2* reference genome, have been remapped on the *NC_000913.3* genome. The same genomic regions are used to train and evaluate all models, at positions 2,738,785, 3,667,115 and 4,131,280 (in accordance to the previous study), as indexed by the *RefSeq* genome (accession: *NC_000913.3*). Both the sense and anti-sense components of these strands are thereby included within the same sets to warrant no unforeseen information transfer between the different sets. The minimum loss on the validation set is used to determine the point at which the network training is halted. Given the prediction task to be a binary classification problem, the cross-entropy loss is used. The performance metrics of the models are obtained by evaluation on the test set.

The Area Under the Receiver Operating Characteristics Curve (ROC AUC) and Area Under the Precision Recall Curve (PR AUC) are the used evaluation metrics. ROC AUC performances represent the area under the curve created by connecting the true positive (y-axis) and false positive (x-axis) rate of the predictions at different thresholds of the output probability. However, due to the imbalance between positive and negative labels, even a small false positive rate can result in set of false positive predictions that heavily outweighs the true positives in absolute count. PR AUC represents the area under the curve created by connecting the precision (precision, y-axis) and recall (x-axis). The impact of false positive predictions for the PR AUC is, unlike the ROC AUC, not proportional to the size of the negative set. Therefore, this metric gives a better depiction of the model’s ability in a practical setting, where the top-k positive predictions are of interest.

### 3.4 Model analysis

Through attention, a score is assigned to a set of upstream hidden states, obtained by the multiplication of the ***q*** and ***k*** vectors and consecutive normalization by the softmax function. For each attention head, this set of scores is investigated in an attempt to determine its function. To obtain a model output at position *n*, 512 (*l*) scores are assigned for each attention head. For a model featuring 36 attention heads and a test set of 1,856,660 positions (samples), a total of about 3.4 · 10^13^ attention scores exists. To reduce the number of values processed, the scores of the upstream hidden states of only a select number of samples is processed. Specifically, in addition to ca. 1% randomly sampled positions of the test set (18,295 samples), we have included the samples with the highest and lowest model output probabilities (500 each), allowing a look into the influence of certain attention heads on the model output and the differences between the profiles of the averaged attention scores.

## 4 Results

### 4.1 The evaluated annotation sets have many different annotated start sites

In order to optimize the set of TSS annotations in *E. coli*, a variety of data sets was first compared. Yan et al. (25) consider TSSs between two sets of annotations to be shared if they are positioned within a distance of five nucleotides. According to this criterion, the distribution of shared TSSs between the four data sets is given in Figure 2a. Only 1,012 positions are shared between all four data sets. The total number of TSSs shared by at least three and two data sets is 2,224 and 5,104, respectively. The fraction of TSSs that are listed by any of the other data sets is the least for RegulonDB (38.17%) and Cappable-seq (28.33%). SMRT-Cappable-seq (89.78%) and SEnd-seq (73.17%) feature the highest fraction of non-unique TSSs (see Table 1). However, as the sizes of the data sets differ, these percentages should be considered with care. For example, Cappable-seq does have the largest absolute number of shared TSSs.

**Figure 2:**
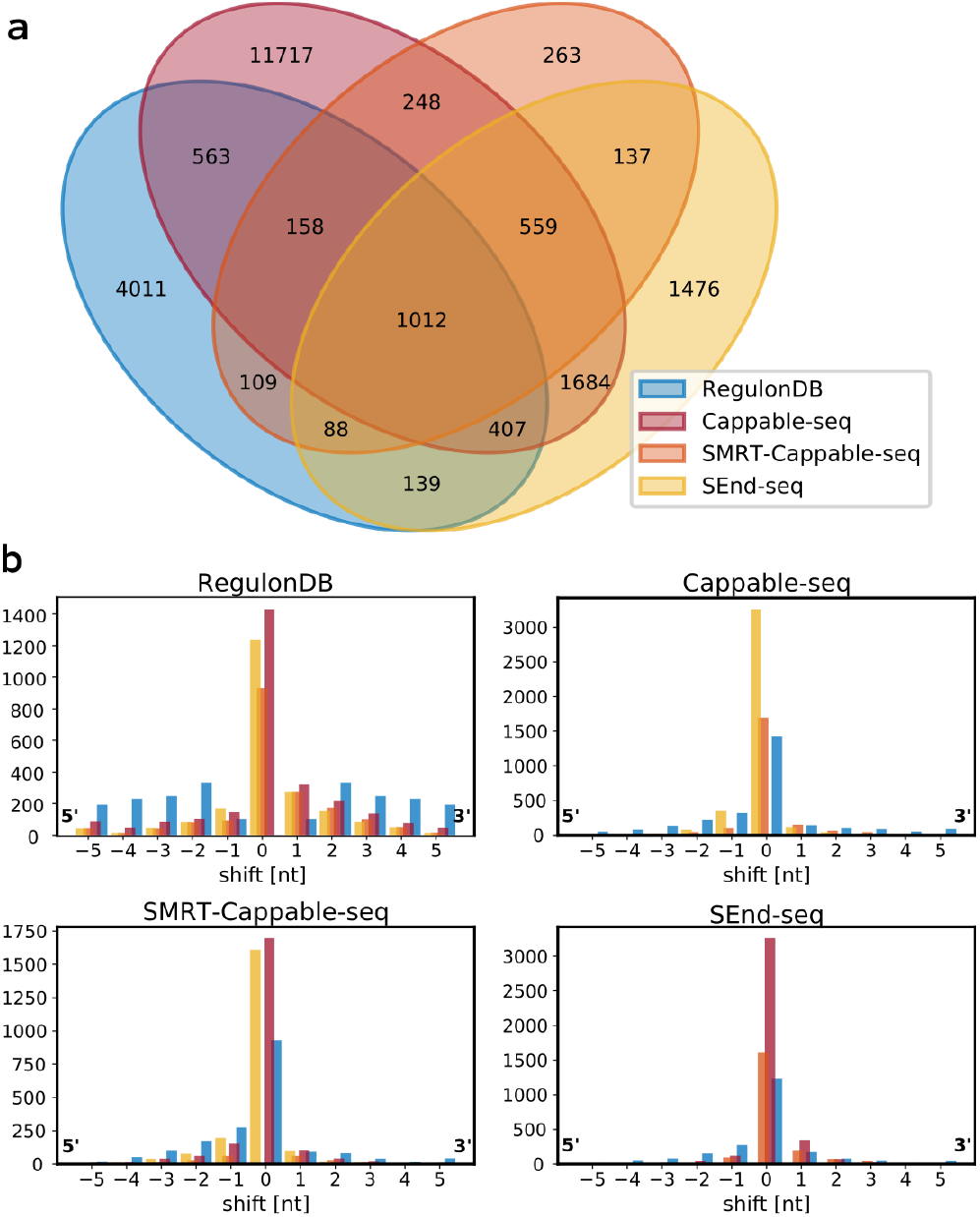
Comparisons between transcription start site (TSS) annotations provided by varying methodologies in *E. coli*. The annotations are obtained from RegulonDB (23) and recent publications introducing novel methodologies from Etwiller et al. (24) (Cappable-seq), Yan et al. (25) (SMRT-Cappable-seq) and Ju et al. (26) (SEnd-seq). (**a**) A venn diagram represents the overlap of annotated TSSs between the different data sources. TSSs are considered to be shared when annotated within 5 nucleotides of one another. (**b**) The distribution of the annotation shift occurring between the shared TSSs within the 5-nucleotide distance used to denote a shared TSS. Distributions are centered on the annotated position for each data set separately.

Figure 2b shows the distributions of the distances between the TSSs within the 5-nucleotide window used in Figure 2a. The percentage of shared TSSs that are annotated at the exact position constitutes only 52.9%, 51.1% and 57.0% for RegulonDB on Cappable-seq, SMRT-Cappable-seq and SEnd-seq, 52.9%, 78.7% and 83.4% for Cappable-seq on RegulonDB, SMRT-Cappable-seq and SEnd-seq, 51.1%, 78.7% and 77.7% for SMRT-Cappable-seq on RegulonDB, Cappable-seq and SEnd-seq, and 57.0%, 83.4% and 77.7% for SEnd-seq on RegulonDB, Cappable-seq and SMRT-Cappable-seq. We can conclude that chosen methods are not able to consistently detect TSSs at single-nucleotide resolution.

Given the differences between annotations present in all data sets, a curated annotation is made that is largely based on the intersection of the discussed datasets rather than the union of the collected annotations. Specifically, the following rules have been applied to include a given TSS to the custom set: (1) the TSS is detected by at least two separate data sets (maximum distance of 5 nucleotides), (2) the exact position of a shared TSSs is determined by the majority vote. In case of a tie, the position is selected based on the novelty of the technique (SEseq > SMRT-Cappable-Seq > cappable-seq > RegulonDB). (3) Finally, given the novelty of SEnd-seq and its high overlap with other data sets, all TSSs detected from this technique have been added. The resulting data set has a total of 6,580 positively labeled positions, denoted as the custom set henceforth.

### 4.2 Improved model performances indicate a better quality of the curated custom set

In order to offer a means of validation about the quality of the annotations and the various steps followed to create the custom set, the performances of different models on the variety of sets have been trained and evaluated. The validation and test sets span identical regions of the genome. Therefore, performances of trained models on the test set can be easily evaluated for different annotations. Table 2 gives a full overview of the performances of all the models on each of the test sets.

**Table 2:** Performance metrics of the transformer network trained and tested on the labeled transcription start sites obtained from RegulonDB (23), Etwiller et al. (24) (Cap(pable)-seq), Yan et al. (25) (SMRT-(Cappable-)seq), Ju et al. (26) (SEnd-seq) and the curated custom set. A model has been trained (rows) and evaluated (columns) on each set of annotations. Both Area Under the Receiver Operating Characteristics Curve (ROC AUC) and Area Under the Precision Recall Curve (PR AUC) are given for each set-up.

| Train set | Test set |  |  |  |  |
| --- | --- | --- | --- | --- | --- |
|  | RegulonDB | Cap-seq | SMRT-seq | SEnd-seq | Custom |
| **ROC AUC** |  |  |  |  |  |
| RegulonDB (23) | **0.882** | 0.815 | 0.923 | 0.882 | 0.885 |
| Cap-seq (24) | 0.790 | **0.961** | 0.938 | 0.945 | 0.945 |
| SMRT-seq (25) | 0.749 | 0.899 | 0.958 | 0.961 | 0.956 |
| SEnd-seq (26) | 0.669 | 0.835 | 0.944 | 0.978 | 0.964 |
| Custom | 0.740 | 0.920 | **0.976** | **0.981** | **0.976** |
| **PR AUC** |  |  |  |  |  |
| RegulonDB (23) | 0.030 | 0.026 | 0.053 | 0.057 | 0.064 |
| Cap-seq (24) | 0.014 | **0.132** | 0.029 | 0.039 | 0.044 |
| SMRT-seq (25) | 0.029 | 0.044 | 0.086 | 0.081 | 0.089 |
| SEnd-seq (26) | 0.035 | 0.052 | 0.098 | **0.128** | 0.137 |
| Custom | **0.039** | 0.057 | **0.141** | **0.128** | **0.141** |

The ROC/PR AUC scores show that the model trained on the custom annotations has the best overall performances on each of the test sets. The highest ROC AUC score on the SMRT-Cappable-seq and SEnd-seq test set is returned by the model trained on the custom set. A possible explanation lies with a lower occurrence of false negative predictions in the model’s output due to a reduced number of missing annotations in the training data. False negative predictions come at a higher cost than false positive predictions, as the true positive rate (y-axis) is more sensitive to change than the false positive rate (x-axis) as an effect of the different sizes of the denominator (i.e. class imbalance).

The low scores of the PR AUC metric illustrate that, despite the high ROC AUC scores, the absolute number of false positive labels outweighs the true positive predictions. Even though these metrics are in line with the state-of-the-art performances previously reported, application of the model on the full genome still results in models with a low precision/recall ratio.

The overall higher performances of the models trained and tested on the custom annotations provide an affirmation that the combined set of transcription start sites, obtained through the steps as outlined in the previous section, have better properties than the individual data sets. As such, all further results are obtained from the model trained and evaluated on the custom set.

### 4.3 The majority of attention heads is highly selective towards certain promoter regions

The attention mechanism is used by the model to selectively collect information from a large pool of data. For each position, the attention head assigns scores to each of the 512 upstream positions. The average and maximum scores at each position (Supplementary Figure S1 and S2) reveal that the majority of attention heads is highly selective about the location from which they extract information. In other words, higher scores are almost exclusively limited to certain promoter regions.

As explained in Material & Methods, the calculation of the attention scores is obtained through the multiplication of the ***q*** and ***k*** vector, each derived from ***h***. The hidden state incorporates information about the nucleotide sequence, its positioning with respect to the other hidden states, and the information added by the residual blocks of previous layers. Based on the averaged attention scores showing a distinct selection as a function of (relative) positioning of the hidden states, it is clear that the model successfully differentiates between the positional and nucleotide information embedded within ***q*** and ***k***. Given the number of samples used, it is inconceivable that the low average and maximum scores along the vast majority of the upstream positions of certain attention heads are solely a result of the sequence information.

### 4.4 Attention heads show to specialize towards detecting transcription factor binding sites

The majority of attention heads that have a high spacial selectivity is delimited in the region spanning from −100 to +20 with respect to the TSS (Supplementary Figure S3 and S4). Note that, through the presence of a convolutional layer in the attention heads, the ***q*** and ***k*** vectors are derived from the seven neighboring hidden states centered at each position (Material & Methods). As such, the information acquired by the attention head at a given position constitutes the 7-mer centered at that position.

For each attention head, sequence motifs were created based on the 50 highest-scoring samples at the position with the highest averaged score (Supplementary Figure S5 and S6). These sequence motifs reflect, in addition to the positional information, the characteristics of the sequence information that result in high attention scores. For example, some of the attention heads in the second layer of the network have an exact overlap with the binding boxes of the RNA polymerase. Here, the 1st and 2nd attention head focus on nucleotide bases ranging from approximately −12 to −6, which interact with region 2 of several *σ* transcription factors. These make up parts of the RNA polymerase holoenzyme as it forms the open complex (32). The motif obtained through selection of the highest scoring sequences is analogous to the reported consensus sequence (TATAAT) of the constitutive *σ*^70^ factor (Figure 3a). The 6th attention head of the 2nd layer overlaps with the −35 box, to which region 4 of the *σ*^70^ transcription factor (33) binds. The obtained sequence motif (Figure 3b) is again analogous to the reported consensus sequence (TTGACA). Nucleotides at the −6 and −5 sites have affinity to *σ*_1.2_ (34), focused on by attention head 2 of layer 1. Feklistov et al. (35) report the presence of GGGA downstream of −10 motif to allow transcription even without the −35 element (Figure 3c). Barne et al. describes the interaction of region *σ*_2.5_ of the RNAP enzyme with the extended −10 promoter region (TGn) (36), spanning the positions −15 to −12. This region is focused on by attention head 4 of layer 2. Alternatively, the positioning and obtained sequence motif (Figure 3d) show resemblance with the −10 element of the *σ*^54^ transcription factor (TTGCAA) (37). The averaged scores and sequence motif of attention head 3 of layer 2 show high similarity with the binding profile of the transcription factor ArcA. There have been 76 binding sites identified in the promoter regions of operons for *E. coli*. The majority of these detected binding sites are situated within the region spanning from −50 to +1. The binding site consensus sequence is the repeat element (TGTTAA) with a distance of 11 base pairs (38). The second and third element (GT) are the most conserved, similar to the constructed sequence motif (Figure 3e). In general, attention heads that do not show to be delimited to certain locations but show strong sequence motifs can constitute transcription factors with varying binding sites within the promoter region. For example, attention head 2 of layer 3 (Figure 3f) returns a motif that almost exactly matches the −35 element of the *σ*^fecI^ RNAP holoenzyme (GGAAAT), but is not specific to the position of this motif with respect to the transcription start site.

**Figure 3:**
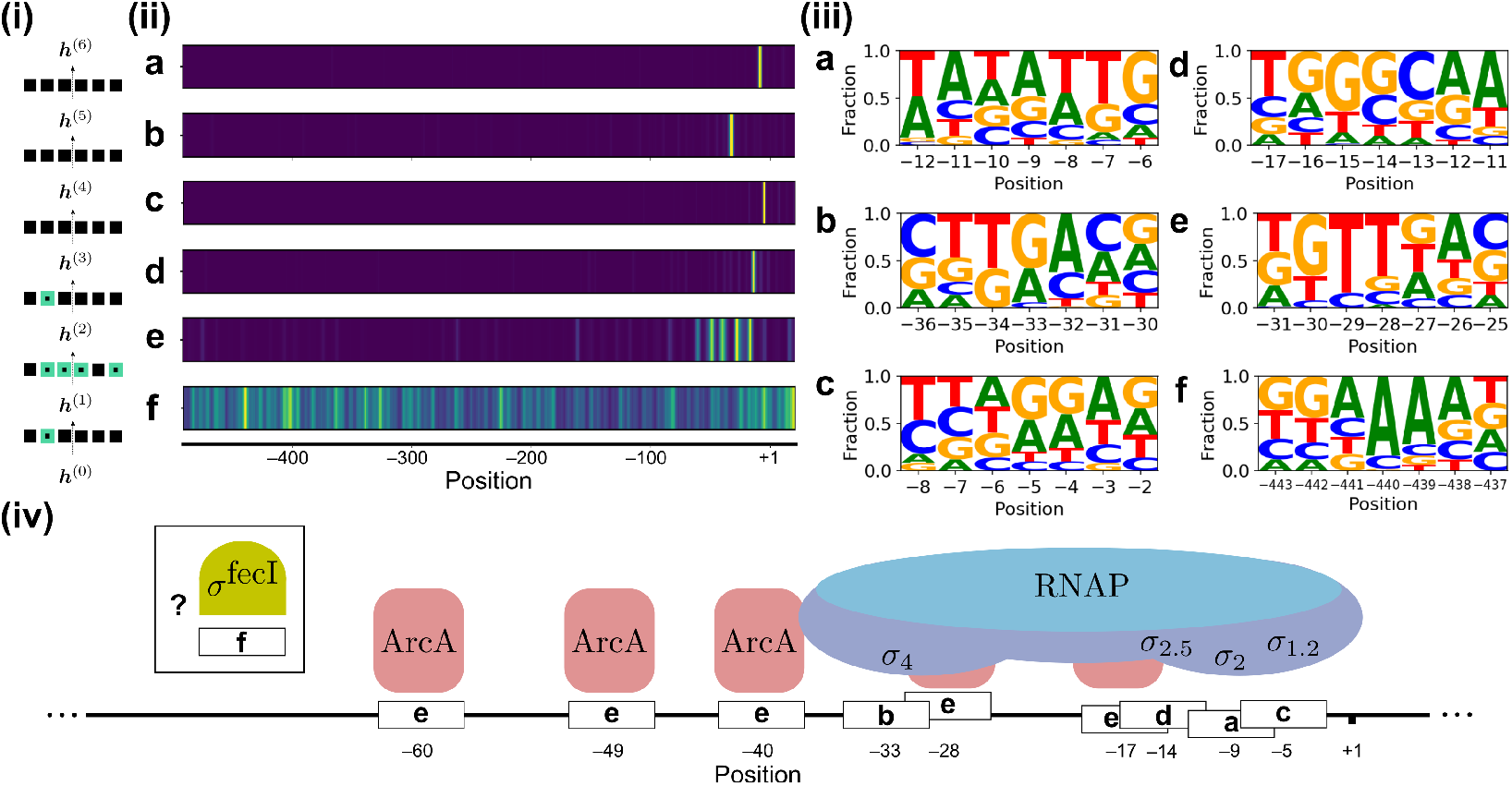
An illustration on some of the information extracted from the trained transformer network based on the analysis of attention heads. **(i)** A depiction of the attention heads (squares) used to calculate the intermediary hidden states ***h***. There are six attention heads for each layer. The green squares show the positions of the attention heads analyzed here within the model architecture. **(ii)** For the six discussed attention heads, the average score along the 512 upstream hidden state are shown. Several attention heads are highly targeted based on positional information, showing high scores in yellow and low in blue. The scores are normalized for each attention head. **(iii)** For every attention head, sequence motifs were made based on the 50 highest scoring samples for the position with the highest average score, illustrating the sequence information that returns high scores for each attention head. **(iv)** Based on the sequence and positional information from (ii) and (iii), a clear match exists that links the specialization of certain attention heads to known factors involved in the transcription process.

Due to the complexity of the subject and the vast amount of literature, it is difficult to verify the function of attention heads without doing extensive research for all potential elements. Additionally, not all mechanisms related to the transcription process are identified. Attention head 1 of layer 6 is highly specific for position −82, which is generally associated with the binding box of th Lac Repressor (39). Alternatively, the −82 site has been reported to overlap with other transcription factor binding sites, such as IscR (40) and the cAMP receptor protein (41).

### 4.5 The influence of the attention heads on the model can be mapped

To further investigate the importance of each attention head with respect to the final model prediction, two strategies were applied. First, the correlation between the attention scores at each position with the model output is quantified using the Spearman’s rank correlation (Supplementary Figure S7). As the hidden state ***h*** at any nucleotide position passes through multiple layers before obtaining a final output prediction (***h***^(0)^ →… → ***h***^(*k*)^), it is expected that the information embedded within the intermediary hidden states gradually reflects the model output. This information is thereby propagated to the attention scores, as ***q*** and ***k*** are derived from ***h***. This can be observed from the maximum and/or minimum correlation coefficient at any position of the attention heads, which increases gradually in the first three layers. The data shows a high correlation coefficient between the attention scores of the third layer and the final model output (~0.85). The majority of the positions has attention scores that are, for most samples, close to zero (e.g. attention head 6 of layer 3). As such, it is likely that the high correlation coefficient of the attention scores of the last layers is an effect of the accumulated information from the previous layers that is stored in ***h***. The results indicate that the attention heads of the first two layers have the largest impact on the model output.

A second strategy compares the distribution of the attention scores of the samples with the highest and lowest model outputs in the test set (Supplementary Figure S3 and S4). Based solely on the scores at the positions with the highest average score for each attention head, we can see that the importance of the attention heads related to the *σ* factor binding boxes show the largest discrepancy between the model outputs that return high and low probabilities.

### 4.6 The output probabilities of the model reflect a correlation between sense and antisense

Unlike previous studies on TSSs (27; 28; 29; 30), the output of the transformer-based model results in a continuous output probability profile along the genome. Evaluation of the output profile has offered further insights into the workings of the model and the related biological process. Similar to previous sections, the results evaluated are obtained from the transformer network trained on the custom annotations.

Increased model outputs are generally concurring on both the forward and reverse strand. Figure 4a shows the median values at each position around the TSSs. This shows the increased output probability on both the sense and antisense strand, where the latter does not only show overall increased activity, but also peaks upstream of the TSSs. Analysis of the probability profile reveals an overall increased output probability for regions with no coding sequence present on either strand. The median output probability over the full region of the test set equals 0.0052. The median probability over all nucleotide positions with a coding sequence present on either strand is 0.0049, a value in line with the median over the full test set. The median value is more than doubled (0.0105) for nucleotides situated in between coding sequences. A higher probability for antisense is not only present for bordering and outward-facing coding sequences, as a higher model output on both strands is prevalent for bordering coding sequences within operons. Using the operon mapping featured on RegulonDB, the median probability within the non-coding sequences of operons is 0.0079 for sense and 0.0071 for antisense. These short sequences have a large number of peaks, where the median over the maxima of each region is ca. ten times the baseline with 0.0415 and 0.0480 for sense and antisense, respectively.

**Figure 4:**
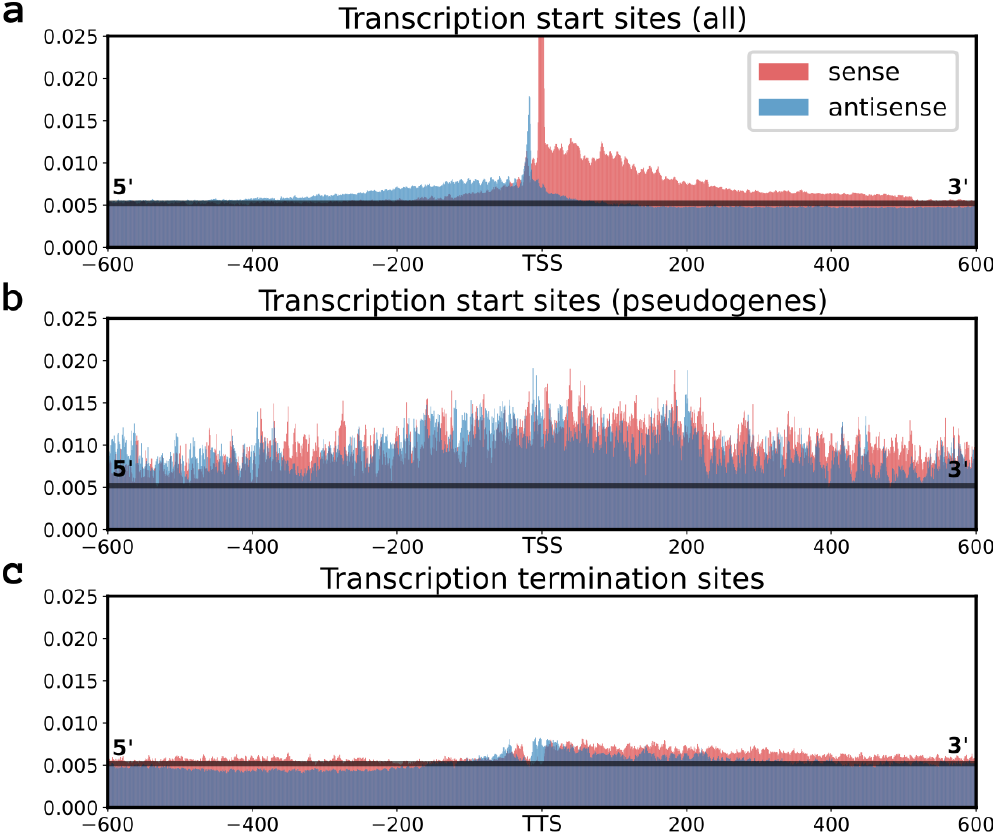
An overview of several plots that describe the prediction characteristics of the model. Data is obtained by the model trained and evaluated on the custom test set. the median output value over the full test set of 0.053 is represented as a horizontal black line in (a,b,c). Each plot gives the median model output for all positions within a 601-nucleotide window centered on the site of interest. Displayed are the median model outputs for the positions around all the (**a**) Transcription Start Sites (TSSs), (**b**) the TSSs of genes annotated as pseudogenes and (**c**) all the transcription termination sites, obtained from Ju et al. (26).

Transcription termination sites were obtained from Ju et al. (26) and shown in Figure 4c. The median scores show a slightly increased probability score around the transcription termination sites for both strands with a shifted dip at the center. Lastly, pseudogenes were found to encompass areas with a substantial increase of output probability along both coding and non-coding nucleotide sequences, resulting in a high number of false positives. The median output probability within the coding sequences of pseudogenes is 0.0111. Most interestingly, the profile around these regions is unique, as no patterns can be discerned. Figure 4b gives the median values for each position around the TSSs of coding sequences annotated as pseudogenes. Supplementary Figure S11c gives an example of a pseudogene region.

Supplementary Figure S11 shows the model output along the custom TSSs annotations as displayed on the UCSC browser (see Supplementary Data). The figure shows examples of the close interaction between the sense and antisense of the model output. The mapped reads of the SMRT-Cappable-seq experiment are also displayed.

## 5 Discussion

The advancements in the field of functional genomics and deep learning offer new opportunities towards gaining understanding of biological processes based on machine learning. Today, existing methods focus on mapping the relation of the input sequence with respect to the model output, and are therefore applicable irrespective of the model architecture (19; 22). However, sensitivity analysis of the input w.r.t. the output can be limiting when the biological process driving the annotation task involves complex mechanisms and/or many factors. These can, for example, involve overlapping binding regions. Moreover, attribution scores on the input sequence are typically averaged for a larger number of inputs, making it hard to detect motifs of importance that are not bound to a single position. To obtain a more layered understanding into the decision-making process of trained models, methods can be created that are bound to the model architecture. As such, it is reasonable to assume that the creation of an explainable deep learning model is a goal that involves both the optimization of the model architecture and the related techniques that seek to explain its functioning. In the case of functional genomics, the training of machine learning models to perform genome annotation tasks generally brings several challenges. These include (i) the imbalance between output classes, where a negative set is presented that is typically several orders of magnitude larger than the positive set, (ii) the lack of high-quality annotations, where a high amount of noise is present, (iii) the dimensions of the input feature space, a multi-million length nucleotide sequence, and (iv) the biological variation of the annotation landscape, which often depends on, among other factors, the (sub)-species, the growth phase, and the growth conditions of the organism.

We recently introduced a custom transformer architecture for processing the full genomic sequence and proved it to be well equipped to address the first and third listed challenge. Next to state-of-the-art performances achieved for all tested annotation tasks (TSSs, translation initiation sites and methylation sites), this framework offers other advantages that improve the efficiency and scalability of the model training process (31). For example, the same set of model weights is applied for calculation of the intermediary states at each position. Thereby, the size of the receptive field (i.e. segment length), the range of input nucleotides that influence the model output, is independent from the number of model weights. Tweaking of the size of the receptive field after training has allowed identification on the influence of excluding information on the model output (31)

After the discovery that attention heads of trained transformer networks are specialized towards mapping semantic relatedness between words (3), we have investigated whether attention heads in the transformer-based model for genome annotation tasks reveal similar characteristics. With over 250 identified potential factors involved in the transcription process of *E. coli* (42), the detection of TSSs serves as a perfect case study to delve into the decision-making process of trained models.

A custom set of TSSs was created and was shown to return better performances than any individual set of annotations available. Potentially, the selection of TSSs from multiple data sets can prevent the model to fit on systematic noise of individual data sets, and make way with missing annotations. Discrepancies occurring between the different annotations might point to an influence of growth conditions on the expression profile. Given that the *in vivo* expression profiles of Cappable-seq, SMRT-Cappable-seq and SEnd-seq were obtained from *E. coli* under optimal growth conditions, the detection of TSSs activated by specialized *σ*-factors, such as *σ*^38^ for stress responses, *σ*^32^ for heat shock response and *σ*^28^ associated with flagellar genes (43), are bound to be limited. We have furthermore observed that the incorporated methods do not provide a precision at single-nucleotide resolution, potentially leading to the multi-nucleotide peaks observed for both the attention heads (Supplementary Figure S3 and S4) and model output (Figure 4). The higher number of shifts in the annotations from RegulonDB indicate this data set to be of lower quality. Today, investigation of TSSs or promoter sequences is generally performed using the annotations from RegulonDB (27; 28; 29; 30), a choice that might not be justified.

The output profile of the model reveals broad areas of increased probabilities for the sense strand around TSSs (Figure 4). These multi-nucleotide peaks might be an effect of noise introduced by annotations that are not precise at single-nucleotide precision (2). Alternatively, it is possible that transcripts with 5’ untranslated regions are not fixed and allow a more flexible start site. To evaluate the impact of these multi-nucleotide peaks on the reported PR AUC scores, a post-processing step that averages the output probabilities within a k-mer, while taking the maximum value for the target label was performed. For a prediction task towards 4-mers on the custom data set, a considerable improvement of the PR AUC is attained (0.29). This score stabilizes when the resolution is further reduced, e.g. 10-mers result in a PR AUC of 0.33.

Our findings establish the specialization of the attention heads, which opens a new branch of opportunities towards explainable deep learning in this field. Focusing on the function of individual attention heads by evaluating the attention scores given to the upstream hidden states, we were able to characterize its functioning by answering a variety of question, such as ‘From which positions within the promoter region does the attention head generally extract its information?’ and ‘What is the motif constructed from the best scoring sequences?’. Based on this information, we identified multiple transcription factor binding sites (Figure 3) and related their importance towards the model output by comparing the scores of the best and least scoring samples for these attention heads (Supplementary Figure S5 and S6). In this study, we have evaluated this based on the position with the highest average attention scores, and might therefore be biased against attention heads that are not highly targeted spatially.

We were able to characterize many important elements related to the biological process the model was trained for. As shown, the effectiveness of the transformer model to specialize certain attention heads towards identifying important promoter regions is owed to the convolutional layer that incorporates information from 7 neighboring hidden states. This length works well for the binding boxes of the *σ*^70^ factor, featuring a motif length of six nucleotides, but would logically not be optimal for transcription factors that involve binding regions that are larger than 7 nucleotides (e.g. LacR, ArcA). Altough many of the characterized attention heads (Supplementary Figure S5 and S6) have not been linked to existing transcription factors, further studies on the topic are likely to uncover more matches between the function of attention heads and and the transcription initiation process.

In addition to characterizing the individual attention heads, which has been the focus in this study, future approaches can be outlined that aim to map the relatedness between attention heads and the model output. Higher order interactions of the attention score profiles between different attention heads might uncover another source of information that offers understanding towards the biological process. Attention heads can, for example, be correlated in order to map transcription factors with binding sites larger than 7 nucleotides. Independent mechanisms for transcription might also be revealed this way. For example, this might be applicable for attention heads picking up alternative *σ* factor binding sites, the −6\−5 motif allowing transcription without the −35 element (35), or the extended −10 element allowing transcription without the −10 element (36). Supplementary Figure S10 shows the correlation matrix between the attention scores of the different attention heads.

Processing of the full the genome is unique for this type of study, as the negative set on which the model is trained is normally subsampled (27; 28; 29; 30). The presence of transcription start sites (and many other genomic sites of interest) for neighboring positions is calculated from largely the same information, yet is expected to return a distinct output profile. The availability of the model predictions along the full genome sequence unlocks a new type of information through which the trained model can be characterized and validated. Specifically, this might result in the identification of model flaws or properties on the transcription process that would not be retrieved otherwise.

The increased probabilities along the sense and antisense of non-coding regions corroborate mechanisms of interference as described in the literature (44). Several types of transcription interference exist that involve convergent promoters: promoter competition, sitting duck, occlusion, collision and roadblock interference. Both sitting duck and roadblock interference involve the occurrence of a RNAP polymerase open complex that is too weak or strong for promoter escape. Promoter competition involves the inhibition of bordering promoters, while occlusion and collision are related to the inference of head-on elongating RNAP. Recently, through the analysis of the SEnd-seq data, the occurrence of head-on interference is described to be an under-appreciated mechanism of transcription regulation (26), an explanation that fits well with aforementioned observations. To further improve model performances on TSSs, possible options include the incorporation of both sense and antisense up- and downstream of the TSSs.

RNAP binding sites that do not result in RNA are likely to contribute to false positive predictions. These include regions where transcription initiation is blocked, such as is the case with head-on interference or with sequences that result in an affinity too high for promoter escape. Head-on interference, among other scenarios, might furthermore result in RNAs that are too small for detection by *in vivo* methodologies. The low transcriptional activity common for pseudogenes (45) could be explained by the interfering affinity of the RNA polymerase for both sense at antisense, as indicated by the probability profile of the model output (Figure 4b and Supplementary Figure 6c). Even though the existence of pseudogenes is not fully understood, the distinct profile created by the model output along the full length of both sense and antisense might well make it possible to detect these solely based on the model output profile. In order to further improve model performances on the detection of TSSs, it is possible that the incorporation of both sense and antisense up- and downstream of the TSSs is required, allowing the model to detect possible factors of interference.

To conclude, the study reveals that the attention mechanism of the transformer-based models offers new opportunities for explainable deep learning in genomics. The availability of both sequence and positional information towards determining attention scores reveals the influence of either factor on the combined result. As such, it is possible to differentiate between motifs that are bound to a single position and those that are not. Combined with the observation that attention heads perform distinct functions, we were able to determine both the motif and position related to several essential factors involved in the process. Additionally, this property might result in correlation studies between the attention scores of different attention heads, potentially unveiling the relatedness between regulatory elements or mechanisms. Using all this information, it is possible that further analysis might be aided by *in vivo* validation methods, thereby fueling the discovery of new mechanisms.

## 6 SUPPLEMENTARY DATA

All models and code used are available on GitHub. https://github.com/jdcla/DNA-transformer. Supplementary File 1 features the full technical details of the transformer network and Supplementary Figures. The annotations and model output are given in Supplementary File 2. All annotations and output of the model trained on the custom set can be browsed on UCSC (https://kermit.ugent.be/files/UCSC/UCSC_browser.html).

