## Supplementary File for "Explainable Transformer Models for Functional Genomics in Prokaryotes"

*Supplementary Information and Figures*
### Explainable Transformer Models for Functional Genomics in Prokaryotes.

Jim Clauwaert [1 a], Gerben Menschaert [2 b], Willem Waegeman [1]

[1] *KERMIT, Department of Data Analysis and Mathematical Modelling, Ghent University, Coupure Links 653, 9000 Gent, Belgium and* [2] *BioBix, Department of Data Analysis and Mathematical Modelling, Ghent University, Coupure Links 653, 9000 Gent, Belgium*

#### 1. Supplementary Information

##### 1.1. Transformer model

Here we describe our transformer network for DNA sequence labeling. In Section 1.2, we adapt the auto-regressive transformer architecture of Dai et al. [1] to DNA sequences. Afterwards, an extension to the calculation of attention is described in Section 1.3.

##### 1.2. Basic model

In essence, the annotation of DNA is a sequence labeling task that has correspondences in natural language processing. The DNA sequence is a data set of $n$ nucleotides, i.e. $\mathbb{X} \in \{\boldsymbol{x}^{(1)}, \boldsymbol{x}^{(2)}, ..., \boldsymbol{x}^{(n)}\}$, where $\boldsymbol{x} \in \{A, C, T, G\}$, the task consists of predicting a label $y \in \{0, 1\}$ for each position $\boldsymbol{x}$, where a positive label denotes the occurrence of an event at that position.

The transformer model processes the genome in sequential segments of $l$ nucleotides. During training, a non-linear transformation function $E$ is optimized that maps the input classes $\{A, C, T, G\}$ to a vector embedding $\boldsymbol{h}$ of length $d_{model}$. for nucleotide $x^{(i)}$ on the genome:

$$\boldsymbol{h} = E(x^{(i)}), \qquad x^{(i)} \in \{A, T, C, G\}, \tag{1}$$

where $\boldsymbol{h} \in \mathbb{R}^{d_{model}}$.

The hidden states of each segment $\boldsymbol{H} \in \mathbb{R}^{l \times d_{model}}$, e.g. $[\boldsymbol{h}^{(1)}, ..., \boldsymbol{h}^{(l)}]$, are processed through $k$ layers. As such, the data propagation through the network for any input $\boldsymbol{x}$ follows multiple transformation: $\boldsymbol{x} \to \boldsymbol{h}^{(0,:)} \to ... \to \boldsymbol{h}^{(k,:)} \to \hat{y}$. A simplistic representation of the data flow is given in Figure 1

Within each layer, multi-head attention is calculated for each hidden state. Next, for each hidden state of $\boldsymbol{h}$, the output of the multi-head attention step ($MultiHead$) is summed with the input, i.e. a residual connection, with the final step being layer normalization. The calculations of the output for all hidden states $\boldsymbol{h}$ in layer $t$ at position $m$ of segment $s$ are performed in parallel:

$$\boldsymbol{h}^{(s,t+1,m)} = LayerNorm(\boldsymbol{h}^{(s,t,m)} + MultiHead(\boldsymbol{H}^{(s,t)})), \tag{2}$$

or

$$\boldsymbol{H}^{(s,t+1)} = LayerNorm(\boldsymbol{H}^{(s,t)} + MultiHead(\boldsymbol{H}^{(s,t)})), \tag{3}$$

where $t \in [0, k[$ and $m \in [1, l]$.

After a forward pass through $k$ layers, a final linear combination reduces the dimension of the output hidden state ($d_{model}$) to the amount of output classes. In this study, only binary classification is performed. A softmax layer is applied before obtaining the prediction value $\hat{y}_i$ for nucleotide $x_i$.

###### 1.2.1. Multi-head attention

The core functionality of the transformer network is the attention head. The attention head evaluates the hidden states in $\boldsymbol{H}$ with one another to obtain an output score $\boldsymbol{z}$. Note, the superscript denoting the layer and segment of the following equations are dropped as identical operations are performed at each layer of each segment.

For each hidden state in the segment, the query ($\boldsymbol{q}$), key ($\boldsymbol{k}$) and value ($\boldsymbol{v}$) vectors are calculated:

$$\boldsymbol{q}, \boldsymbol{k}, \boldsymbol{v} = \boldsymbol{h}\boldsymbol{W}^{\boldsymbol{q}}, \boldsymbol{h}\boldsymbol{W}^{\boldsymbol{k}}, \boldsymbol{h}\boldsymbol{W}^{\boldsymbol{v}}, \tag{4}$$

where $\boldsymbol{W}^{\boldsymbol{q}}, \boldsymbol{W}^{\boldsymbol{k}}, \boldsymbol{W}^{\boldsymbol{v}} \in \mathbb{R}^{d_{heads} \times d_{model}}$ and $\boldsymbol{q}, \boldsymbol{k}, \boldsymbol{v} \in \mathbb{R}^{d_{heads}}$. The $\boldsymbol{q}$ and $\boldsymbol{k}$ vectors are used to obtain a score between two hidden states, expressing their relevance with one another in regard to the information represented by $\boldsymbol{v}$.

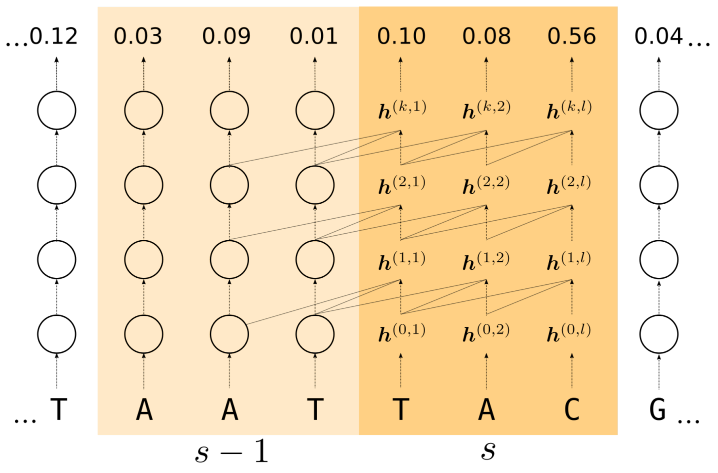


Figure 1: Illustration of the connectivity of intermediary values of the transformer architecture. The full genome is processed in sequential segments $s$ with length $l$. First, the input nucleotide is transformed into a vector embedding $\boldsymbol{h}^{(0)}$, after which it is processed by $k$ consecutive residual blocks ($\boldsymbol{h}^{(0)} \rightarrow ... \rightarrow \boldsymbol{h}^{(k)}$). The output probability is obtained by sending the final hidden state $\boldsymbol{h}^{(k)}$ through a set of fully-connected layers. For the calculation at each residual block, the last $l$ hidden states of the previous layer are applied. For example, $\boldsymbol{h}^{(1,l)}$ is calculated by using the hidden states $[\boldsymbol{h}^{(0,1)}, ..., \boldsymbol{h}^{(0,l)}]$. Hidden states from the previous segment $(s-1)$ are made accessible for the calculation of the hidden states in segment $s$.

For each hidden state at position $m$ of the segment, the attention score $\boldsymbol{z}$ is calculated by evaluation of its vector $\boldsymbol{q}$ with the $\boldsymbol{k}$ and $\boldsymbol{v}$ vectors derived from the other hidden states in the segment:

$$\boldsymbol{z}^{(m)} = softmax(\sum_{i=1}^{l} \frac{\boldsymbol{q}^{(m)} \cdot \boldsymbol{k}^{(i)}}{\sqrt{d_{heads}}}) \cdot \boldsymbol{v}^{(i)}. \tag{5}$$

The $softmax$ function is used to rescale the weights assigned to the vectors $\boldsymbol{v}$ to sum to 1. Division by the square root of $d_{heads}$ is applied to stabilize gradients [2].

The calculation of attention within the attention head is performed in parallel for all hidden states in $\boldsymbol{H}$:

$$\boldsymbol{Q}, \boldsymbol{K}, \boldsymbol{V} = \boldsymbol{H}\boldsymbol{W}^{\boldsymbol{q}\top}, \boldsymbol{H}\boldsymbol{W}^{\boldsymbol{k}\top}, \boldsymbol{H}\boldsymbol{W}^{\boldsymbol{v}\top}, \tag{6}$$

$$\begin{aligned} \boldsymbol{Z} =& Attention(\boldsymbol{H}), \\ =& softmax(\frac{\boldsymbol{Q}\boldsymbol{K}^\top}{\sqrt{d_{heads}}})\boldsymbol{V}, \end{aligned} \tag{7}$$

where $\boldsymbol{Q}, \boldsymbol{K}, \boldsymbol{V} \in \mathbb{R}^{l \times d_{heads}}$ and $\boldsymbol{Z} \in \mathbb{R}^{l \times d_{heads}}$. Here, the $softmax$ function is applied to every row of $\boldsymbol{Q}\boldsymbol{K}^\top$.

To increase the capacity of the model, the input is processed by multiple attention heads ($n_{heads}$) present within each layer, each featuring a unique set of weight matrices $\boldsymbol{W}^{\boldsymbol{q}}$, $\boldsymbol{W}^{\boldsymbol{k}}$, $\boldsymbol{W}^{\boldsymbol{v}}$—optimized during training. Having multiple sets of $\boldsymbol{W}^{\boldsymbol{k}}$, $\boldsymbol{W}^{\boldsymbol{q}}$ and $\boldsymbol{W}^{\boldsymbol{v}}$ allows the model to extract multiple types of information from the hidden states.

The output of the multi-head attention unit is obtained by concatenation of all $\boldsymbol{Z}$ matrices along the second dimension and multiplication by $\boldsymbol{W}^{\boldsymbol{m}}$. This creates an output with dimensions equal to $\boldsymbol{H}$:

$$MultiHead(\boldsymbol{H}) = ColConcat(Z^{(1)}(\boldsymbol{H}), ..., Z^{(n_{heads})}(\boldsymbol{H}))\boldsymbol{W}^{\boldsymbol{m}} \tag{8}$$

where $\boldsymbol{W}^{\boldsymbol{m}} \in \mathbb{R}^{n_{heads}d_{heads} \times d_{model}}$.

##### 1.2.2. Recurrence

To process the full genome sequence, a recurrence mechanism is applied, as described by Dai et al.[1]. This allows for the processing of a single input (i.e. the genome) in sequential segments of length $l$. In contrast with calculation of the attention heads described in the previous section, only upstream hidden states are used to calculate the output of $\boldsymbol{h}$. In each layer, hidden states $[\boldsymbol{h}^{(m+1)}, ..., \boldsymbol{h}^{(l)}]$ are masked when processing $\boldsymbol{z}^{(m)}$, $m \in [1, l]$.

In order to extend the receptive field of information available past one segment, hidden states of the previous segment $s-1$ are accessible for the calculation of $\boldsymbol{h}^{(s,t+1)}$. The segment length $l$ denotes the span of hidden states used to calculate attention. Therefore, $\boldsymbol{H}^{(s,t,m)}$, representing the collection of hidden states used for the calculation of multi-head attention at position $m$ in layer $t+1$ of segment $s$, consists of $l$ hidden states typically spanning over segment $s$ and $s-1$:

$$\boldsymbol{H}^{s,t,m} = [SG(\mathbf{h}^{s-1,t,m+1} \quad ... \quad \mathbf{h}^{s-1,t,l}) \quad \mathbf{h}^{s,t,1} \quad ... \quad \mathbf{h}^{s,t,m}]. \tag{9}$$

$SG$ denotes the stop-gradient, signifying that during training, no weight updates of the model are performed based on the partial derivatives of given hidden states with the loss. This alleviates training times, as full backpropagation through intermediary values would require the model to retain the hidden states from as many segments as there are layers present in the model, a process that quickly becomes unfeasible for a model with a large segment length or high amount of layers. Figure 1 gives an overview of the model architecture adopting the recurrence mechanism.

##### 1.2.3. Relative Positional Encodings

Next to the information content of the input, positional information of the hidden states is relevant towards the calculation of attention. Unlike the majority of other machine learning methods in the field (e.g. linear regression, convolutional/recurrent neural networks), the architecture of the model does not inherently incorporate the relative positioning of the inputs with respect to the output. Positional information is added through the use of positional embeddings. These introduce a bias during calculation of $\boldsymbol{z}$ that is related to the vector representation and relative distance of the evaluated hidden states. Predefined biases are learned during the training phase and have been found to work well on test data. Attention between positions $m$ and $o$, in function of the hidden state in $m$, is evaluated by expanding the algorithm [1]:

$$
\begin{aligned}
A(\boldsymbol{H})_{m,o} = \underbrace{\boldsymbol{H}_{m,:}\boldsymbol{W}^{\boldsymbol{q}\top}\boldsymbol{W}^{\boldsymbol{k},\boldsymbol{H}}\boldsymbol{H}^{\top}_{o,:}}_{(a)} + \underbrace{\boldsymbol{H}_{m,:}\boldsymbol{W}^{\boldsymbol{q}\top}\boldsymbol{W}^{\boldsymbol{k},\boldsymbol{R}}\boldsymbol{R}^{\top}_{m-o,:}}_{(b)} + \\
\underbrace{\boldsymbol{u}^{\top}\boldsymbol{W}^{\boldsymbol{k},\boldsymbol{H}}\boldsymbol{H}^{\top}_{o,:}}_{(c)} + \underbrace{\boldsymbol{v}^{\top}\boldsymbol{W}^{\boldsymbol{k},\boldsymbol{R}}\boldsymbol{R}^{\top}_{m-o,:}}_{(d)},
\end{aligned} \tag{10}
$$

$$
\boldsymbol{Z} = softmax(\frac{A(\boldsymbol{H})}{\sqrt{d_{heads}}})\boldsymbol{V}, \tag{11}
$$

where $\boldsymbol{W}^{\boldsymbol{k},\boldsymbol{H}}$ (or $\boldsymbol{W}^{\boldsymbol{k}}$) is the weight matrix used to calculate the key (K) matrix for the input hidden states. $\boldsymbol{W}^{\boldsymbol{k},\boldsymbol{R}} \in \mathbb{R}^{d_{heads} \times d_{model}}$ is a new weight matrix used in relation to positional information embedded in $\boldsymbol{R}$. $\boldsymbol{R} \in \mathbb{R}^{l \times d_{model}}$ is a matrix defining biases related to the distance between $m$ and $o$. $\boldsymbol{u}, \boldsymbol{v} \in \mathbb{R}^{d_{heads}}$ are vectors that relate to content and positional information on a global level, optimized during training. Several elements make up the algorithm to obtain $A(\boldsymbol{H})$:

(**a**) Attention based on query and key values of the input hidden states, described in the previous section.

(**b**) Bias based on hidden state in $m$ ($\boldsymbol{H}_{m,:}\boldsymbol{W}^{\boldsymbol{q}\top}$) and distance to $o$ ($\boldsymbol{W}^{\boldsymbol{k},\boldsymbol{R}}\boldsymbol{R}^{\top}_{m-o,:}$).

(**c**) Bias based on hidden state in $o$ ($\boldsymbol{W}^{\boldsymbol{k},\boldsymbol{H}}\boldsymbol{H}^{\top}_{o,:}$), unrelated to the relative position to $m$. $\boldsymbol{u}$ is optimized during training.

(**d**) Bias based on distance between $m$ and $o$ ($\boldsymbol{W}^{\boldsymbol{k},\boldsymbol{R}}\boldsymbol{R}^{\top}_{m-o,:}$), unrelated to the content of the hidden states. $\boldsymbol{v}$ is optimized during training.

#### 1.3. Extension: Convolution over Q, K and V

Important differences exist between the input sequence of the genome and typical natural language processing tasks. The genome constitutes a very long sentence, showing low contextual complexity at input level. Indeed, only four input classes exist. Attention is calculated based on the individual hidden states $\boldsymbol{h}$. For example, in the first layer, hidden states of the segment solely contain information on the nucleotide classes. In previous studies, meaningful sites and regions of interest on the genome are specified by (sets of) motifs from neighboring nucleotides.

To expand the information contained in $\boldsymbol{q}$, $\boldsymbol{k}$ and $\boldsymbol{v}$ to represent k-mers rather than single nucleotides, a 1D convolutional layer is implemented that convolves over the $\boldsymbol{q}$, $\boldsymbol{k}$ and $\boldsymbol{v}$ vectors derived from neighboring hidden states, present as adjoining rows in $\boldsymbol{Q}$, $\boldsymbol{K}$ and $\boldsymbol{V}$. The length of the motif, k-mer or kernel is denoted by $d_{conv}$.

To ensure the dimensions of $\boldsymbol{q}$, $\boldsymbol{k}$ and $\boldsymbol{v}$ to remain identical after the convolutional step, as many sets of weight kernels are trained as $d_{heads}$. Furthermore, through padding, the size of the first dimension of the matrices $\boldsymbol{Q}$, $\boldsymbol{K}$ and $\boldsymbol{V}$ can be kept constant. Applied on $\boldsymbol{q}$ we get:

$$
\begin{aligned}
\boldsymbol{q}^{(m,conv)}_{c} = \sum_{i=1}^{d_{conv}} \sum_{j=1}^{d_{heads}} q_j^{(f(m,i))} W^{conv,q}_{i,j,c}, \\
f(m,i) = m - \lceil \frac{d_{conv}}{2} \rceil + i.
\end{aligned} \tag{12}
$$

where $c \in [1, d_{heads}]$ and $\boldsymbol{W}^{\boldsymbol{conv},\boldsymbol{q}} \in \mathbb{R}^{d_{heads} \times d_{conv} \times d_{heads}}$. $\boldsymbol{W}^{\boldsymbol{conv},\boldsymbol{q}}$ is the tensor of weights used to convolve $\boldsymbol{q}$. Applied on the $\boldsymbol{Q}$ matrix the operation is represented as:

$$
\boldsymbol{Q}^{\boldsymbol{conv}}_{m} = \sum_{i=1}^{d_{conv}} \sum_{j=1}^{d_{heads}} Q_{f(m,i),j} W^{conv,q}_{i,j}. \tag{13}
$$

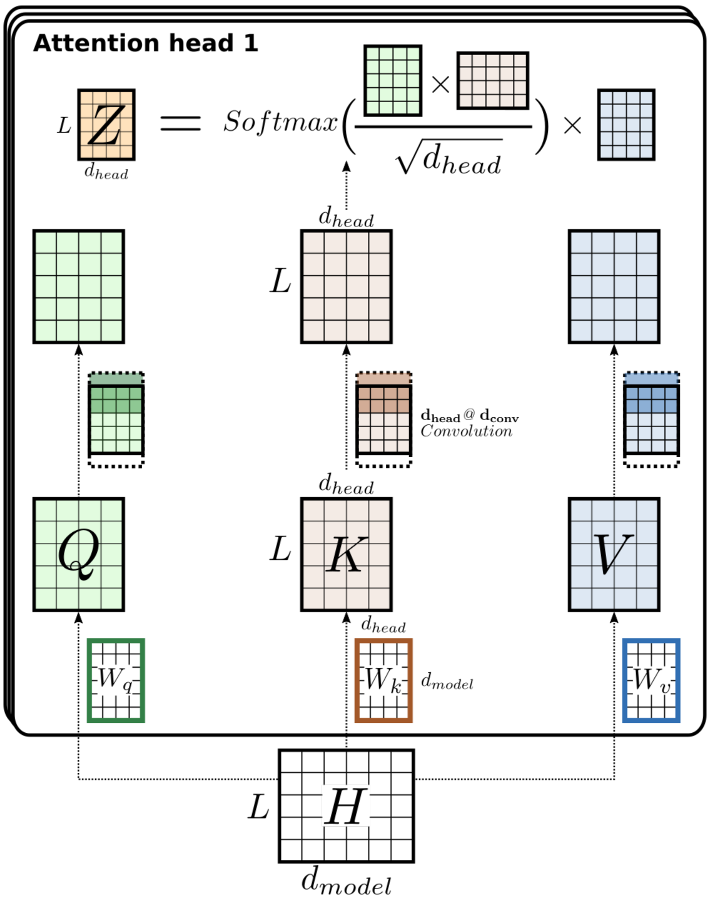


Figure 2: An overview of mathematical operations performed by the attention head to calculate attention $\boldsymbol{z}$ for each hidden state $\boldsymbol{h}$. Operations to calculate attention are performed in parallel for $l$ hidden states ($\boldsymbol{H} \rightarrow \boldsymbol{Z}$). The $\boldsymbol{q}$, $\boldsymbol{k}$ and $\boldsymbol{v}$ vectors are obtained through matrix multiplication of $\boldsymbol{H}$ with $\boldsymbol{W^q}$, $\boldsymbol{W^k}$ and $\boldsymbol{W^v}$, resulting in $\boldsymbol{Q}$, $\boldsymbol{K}$ and $\boldsymbol{V}$. A single convolutional layer using as many kernels as $d_{heads}$ results in the transformation of the individual $\boldsymbol{q}$, $\boldsymbol{k}$ and $\boldsymbol{v}$ vector representations of each input to be derived from the $\boldsymbol{q}$ $\boldsymbol{k}$ and $\boldsymbol{v}$ vectors of $d_{conv}$ bordering nucleotides. Attention $\boldsymbol{Z}$ is thereafter calculated. The schema is kept simple for better understanding and does not include the relative position encodings added to the $\boldsymbol{Q}$ and $\boldsymbol{K}$ matrix, nor does it incorporate the recurrence mechanism.

A unique set of weights is optimized to calculate $\boldsymbol{Q^{conv}}$, $\boldsymbol{K^{conv}}$ and $\boldsymbol{V^{conv}}$ for each layer. To reduce the total amount of parameter weights of the model, identical weights are used to convolve $\boldsymbol{Q}$, $\boldsymbol{K}$ and $\boldsymbol{V}$ for all attention heads in the multi-head attention module. Figure 2 gives a visualization of the intermediate results and mathematical steps performed to calculate attention within the attention head of the extended model.

##### 1.4. Model hyperparameters

The model hyperparameters have been discussed in previous work and have not been altered (Table 1) [3].

Table 1: Overview of the hyperparameters that define the model architecture

| Hyperparameter | Symbol | Value | Hyperparameter | Symbol | Value |
|---|---|---|---|---|---|
| layers | $k$ | 6 | segment length | $l$ | 512 |
| dim. head | $d_{heads}$ | 6 | dim. model | $d_{model}$ | 32 |
| heads in layer | $n_{heads}$ | 6 | conv. kernel size | $d_{conv}$ | 7 |
| learning rate | $lr$ | 0.0002 | batch size | $bs$ | 10 |

#### 2. Code

Code used to train the models discussed in the article are available on GitHub (`https://github.com/jdcla/DNA-transformer`). The code is forked from the GitHub repository of the transformer-XL architecture (`https://github.com/kimiyoung/transformer-xl`).

#### 3. Model output

The model output on the test set region alongside all sets of annotations has been visualized in the UCSC browser and can be accessed here: `https://kermit.ugent.be/files/UCSC/UCSC_browser.html`

#### 4. Supplementary Figures

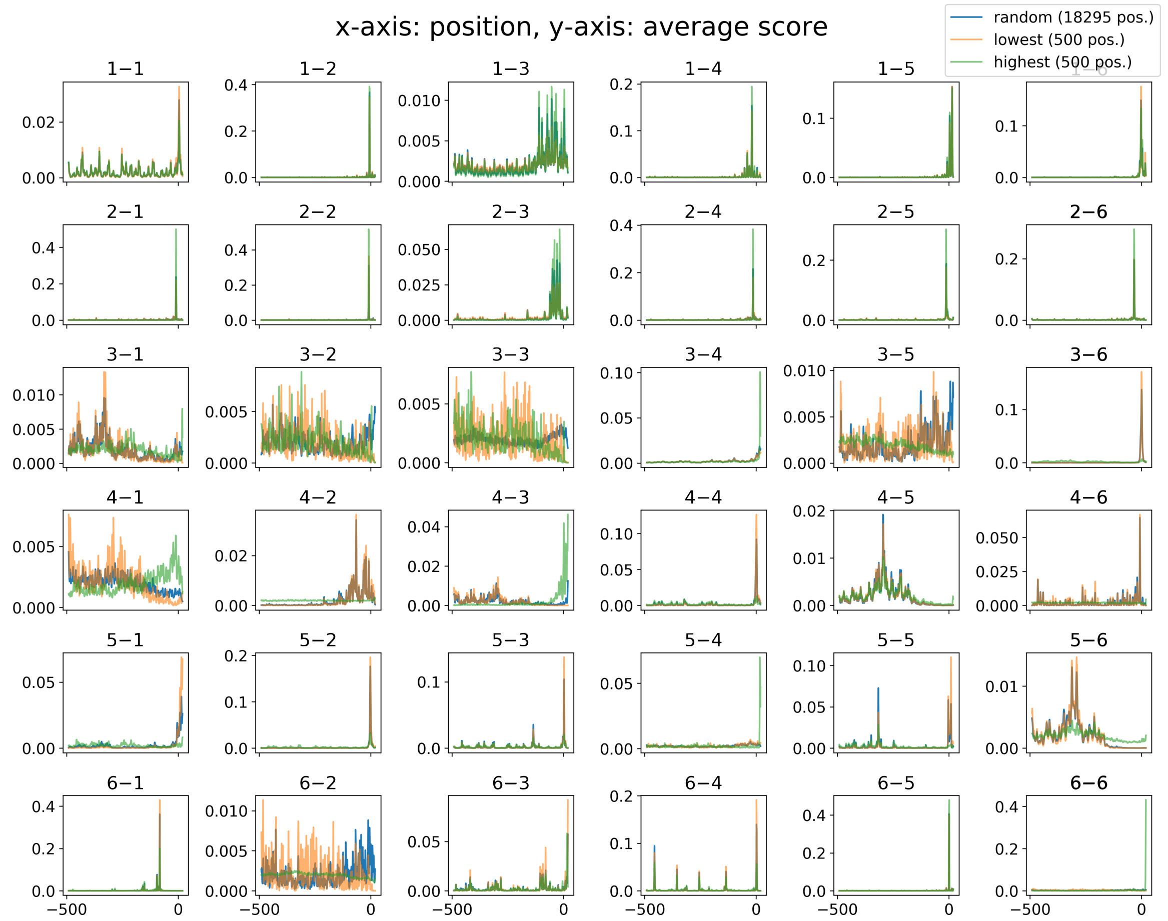


Figure S1: The average weights assigned to each upstream position for the calculation of attention for each of the attention heads (see Equation 5). Attention heads are grouped by layer (first index). Weights are obtained from the model trained and the custom set of annotations. The average score for three sets of samples are compared: a subsample of the full test set (random), accounting for 18,295 samples, and the samples with the 500 highest and lowest model outputs. Weights are obtained from the model trained on the custom set of annotations. Based

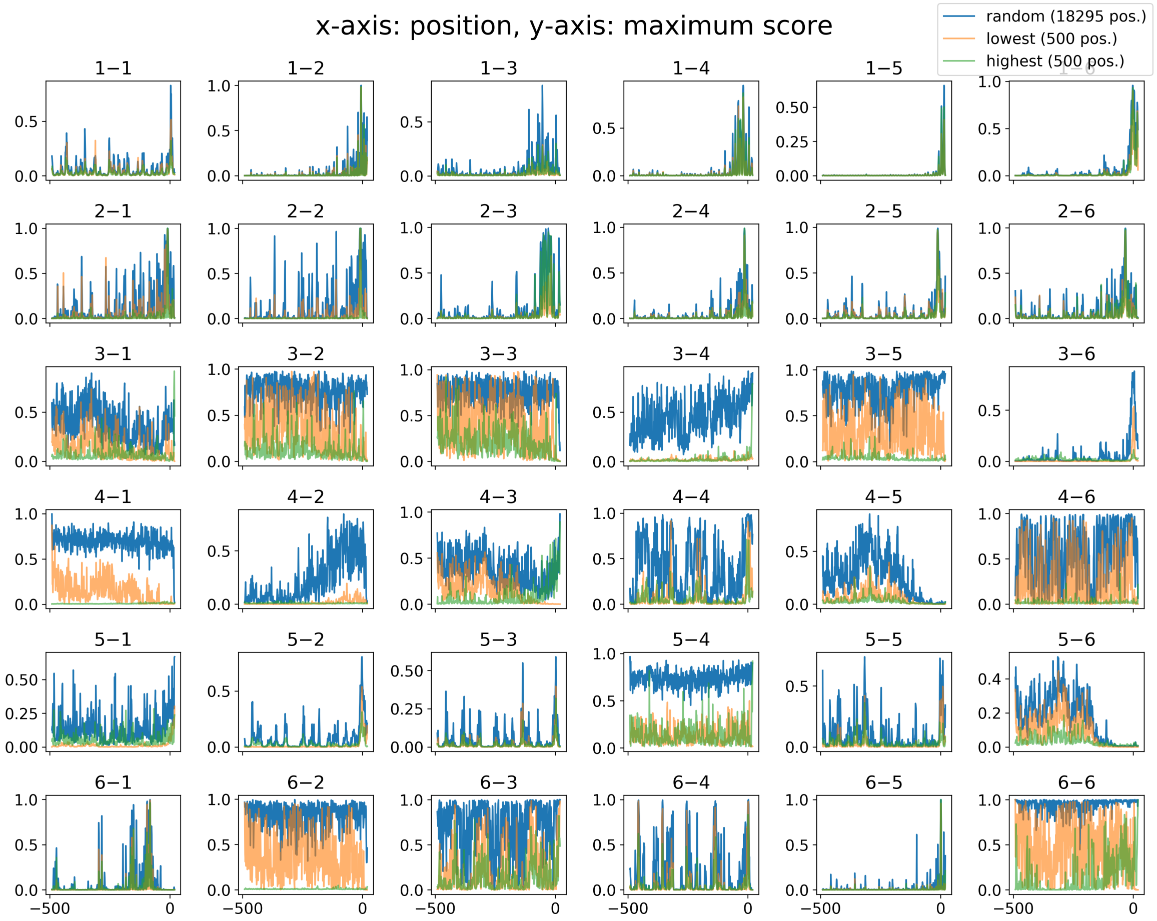


Figure S2: The maximum weights assigned to each upstream position for the calculation of attention for each of the attention heads (see Equation 5). Attention heads are grouped by layer (first index). Weights are obtained from the model trained and the custom set of annotations.The maximum score for three sets of samples are compared: a subsample of the full test set (random), accounting for 18,295 samples, and the samples with the 500 highest and lowest model outputs. Weights are obtained from the model trained on the custom set of annotations.

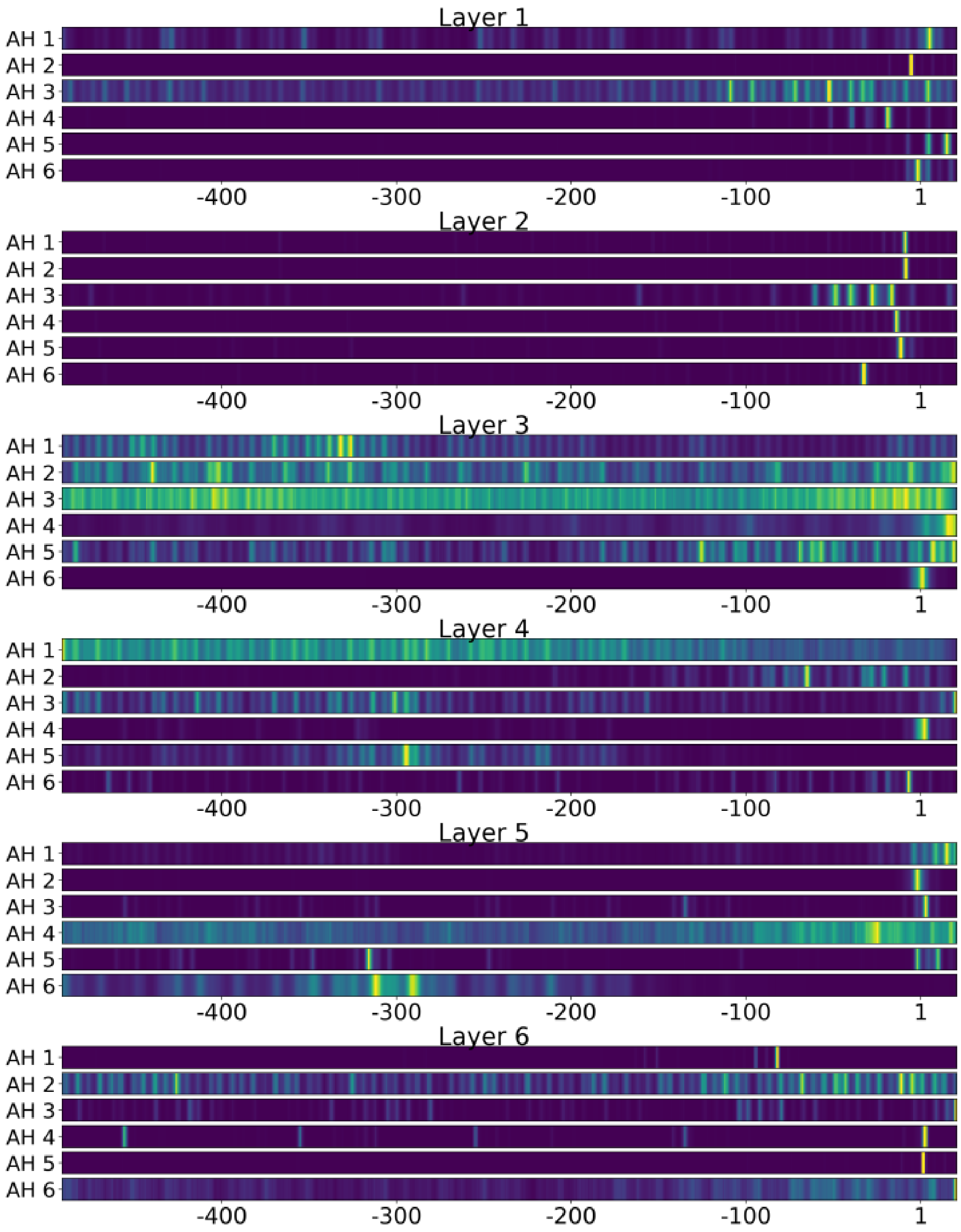


Figure S3: An alternative visualization of the average weights as portrayed by Supplementary Figure S1 for the subsampled test set (18,295 samples). The colormap represents low values as darker blue and higher values as yellow, and are normalized to range from 0 to 1 for each attention head (AH). The weight at each position is calculated based on the 7 neighboring hidden states centered at that position (see Section 1.3).

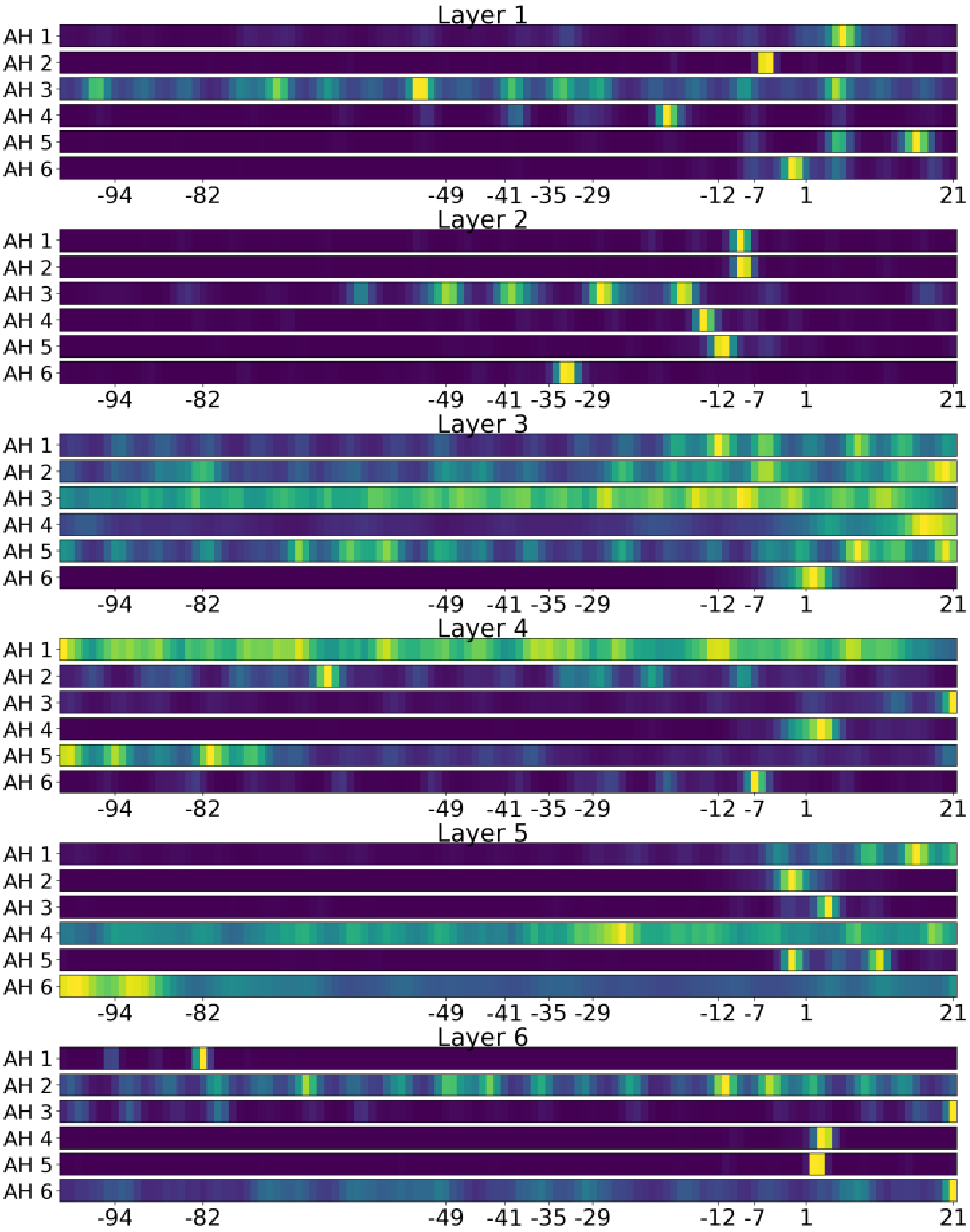


Figure S4: An alternative visualization of the average weights as portrayed by Supplementary Figure S1 for the subsampled test set (18,295 samples), centered on the last 120 hidden states. The colormap represents low values as darker blue and higher values as yellow, and are normalized to range from 0 to 1 for each attention head (AH). The weight at each position is calculated based on the 7 neighboring hidden states centered at that position (see Section 1.3).

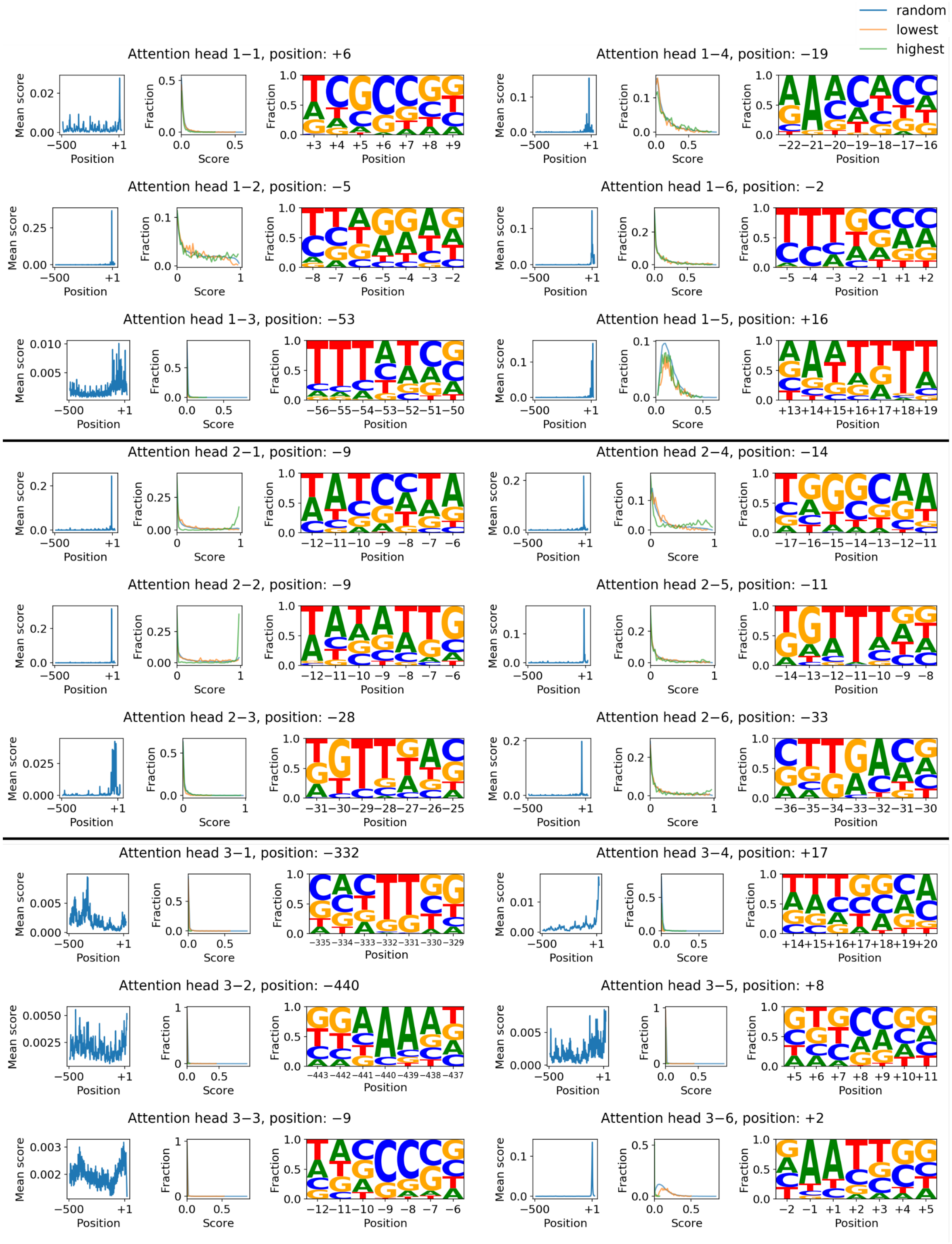


Figure S5: For each of the attention heads, a motif is constructed at the position with the highest average score for all samples (see Supplementary Figure S1 and S3). For each attention head is given: (left) The average scores/weights given to the 512 upstream hidden states, (middle) the distribution of the scores at the position with the highest average score, in addition to the distributions at that position for the sample sets containing the 500 highest and lowest model outputs, (right) a position frequency matrix of the position constructed using the highest scoring samples for that attention head (top 50). Attention heads are grouped by layer (first index). The motifs for the last three layers are given in Supplementary Figure S6.

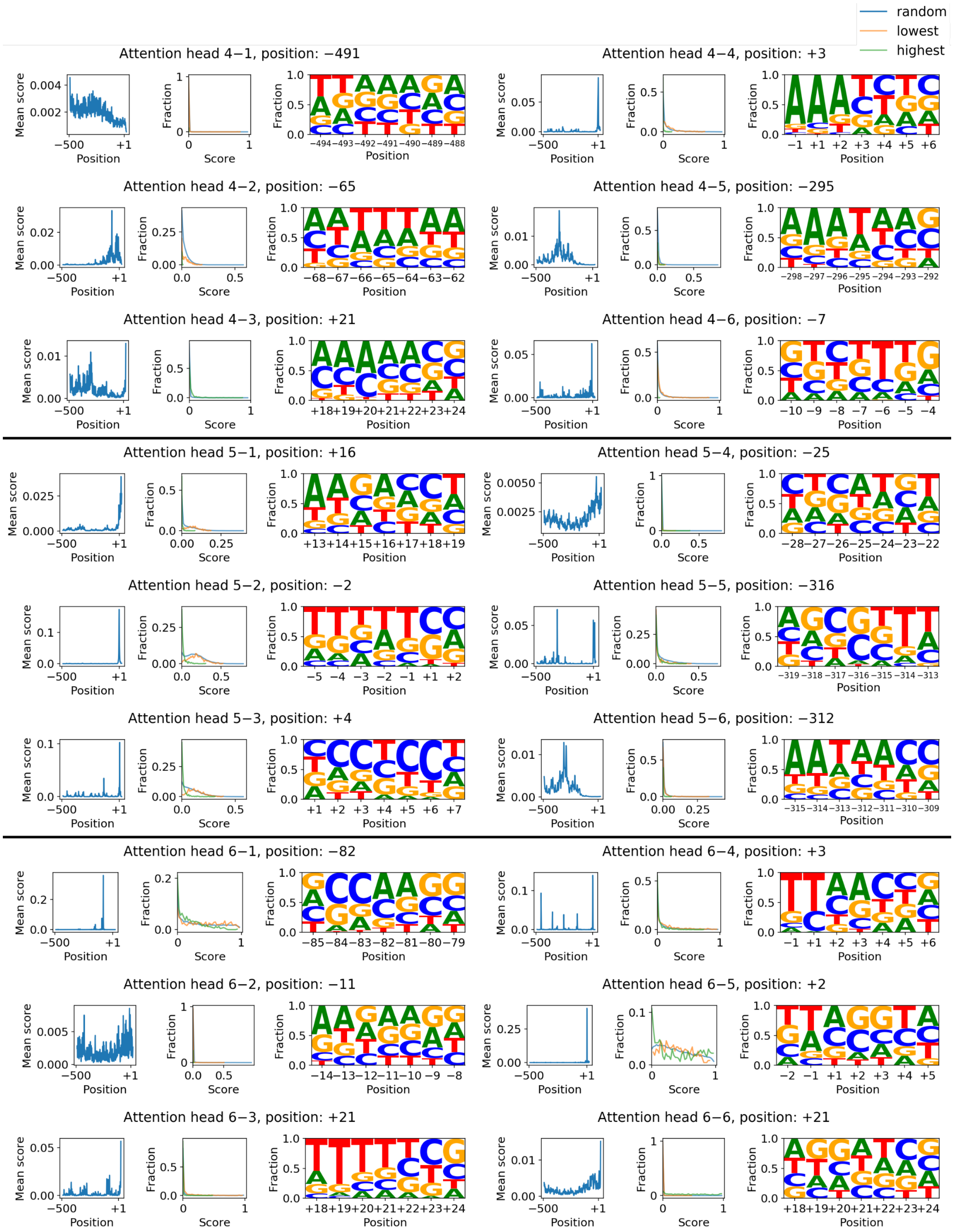


Figure S6: For each of the attention heads, a motif is constructed at the position with the highest average score for all samples (Supplementary Figure S1 and S3). For each attention head is given: (left) The average scores/weights given to the 512 upstream hidden states, (middle) the distribution of the scores at the position with the highest average score, in addition to the distributions at that position for the sample sets containing the 500 highest and lowest model outputs in the test set, (right) a position frequency matrix of the position constructed using the highest scoring weights at that position (top 50). Attention heads are grouped by layer (first index). The motifs for the first three layers are given in Supplementary Figure S5.

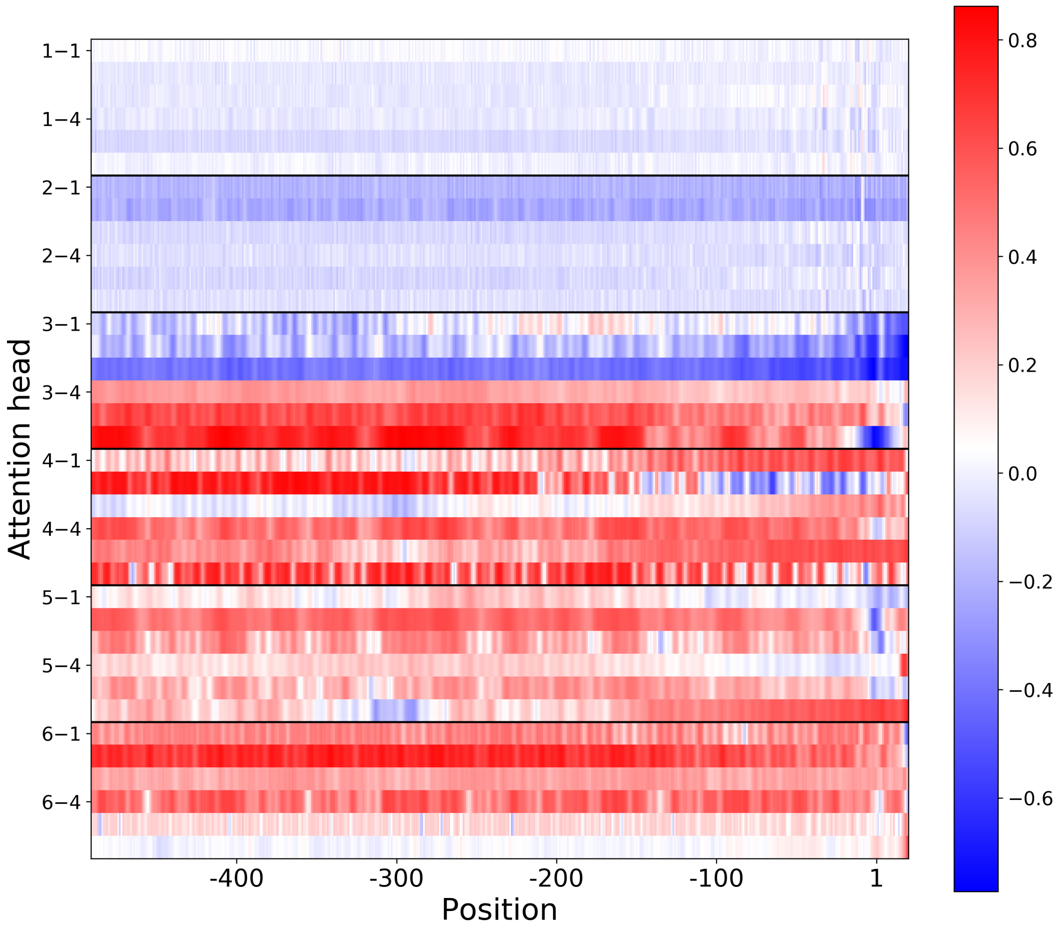


Figure S7: For each of the attention heads, the Spearman's Rank correlation between the score assigned to each (upstream) position and the model output is displayed. Attention heads are grouped by layer (first index).

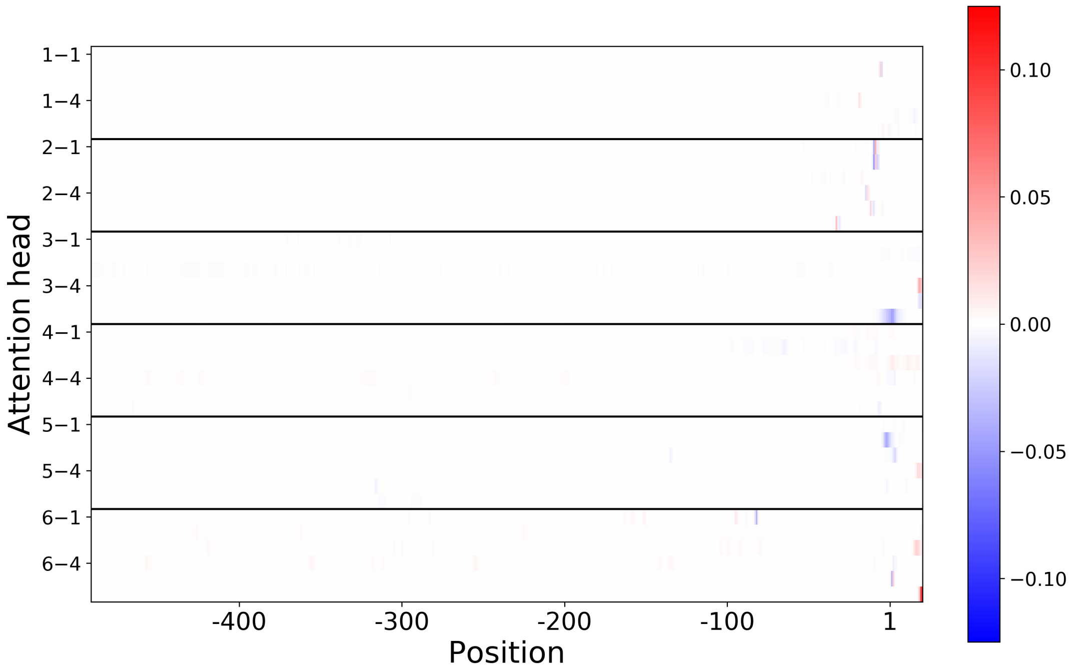


Figure S8: Similar to Supplementary Figure S7, for each of the attention heads, the Spearman's Rank correlation between the score assigned to each (upstream) position and the model output is displayed. Each correlation coefficient has been multiplied by the average weight attributed at that position (Supplementary Figure S1). Attention heads are grouped by layer (first index).

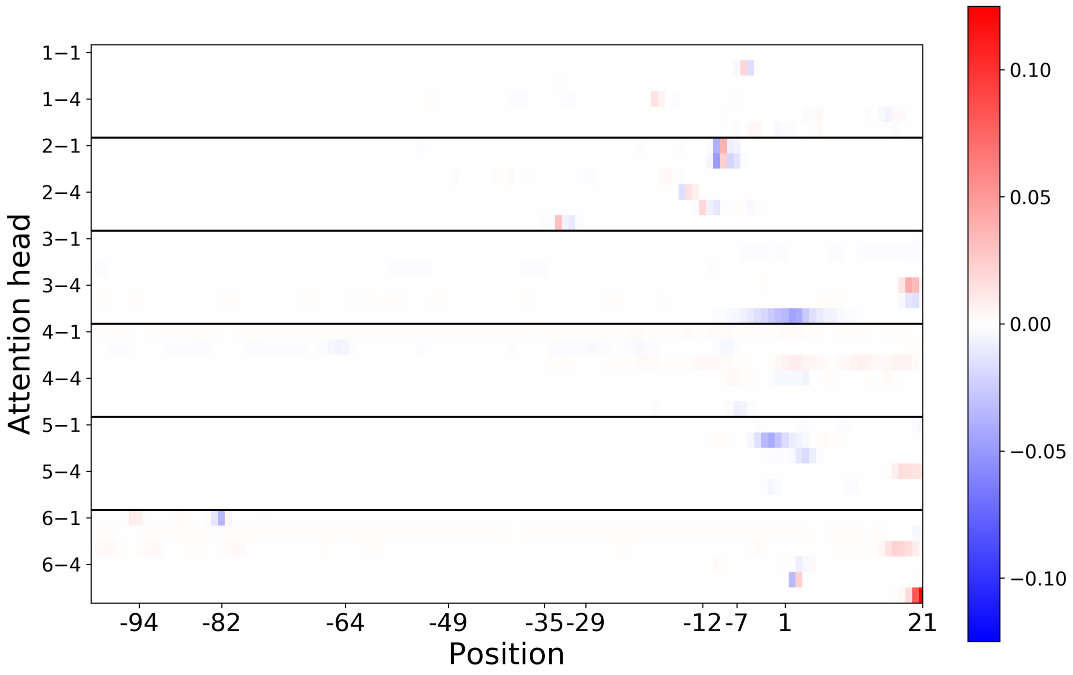


Figure S9: Similar to Supplementary Figure S7, for each of the attention heads, the Spearman's Rank correlation between the score assigned to each of the 120 (upstream) positions and the model output is displayed. Each correlation coefficient has been multiplied by the average score attributed at that position (Supplementary Figure S1). Attention heads are grouped by layer (first index).

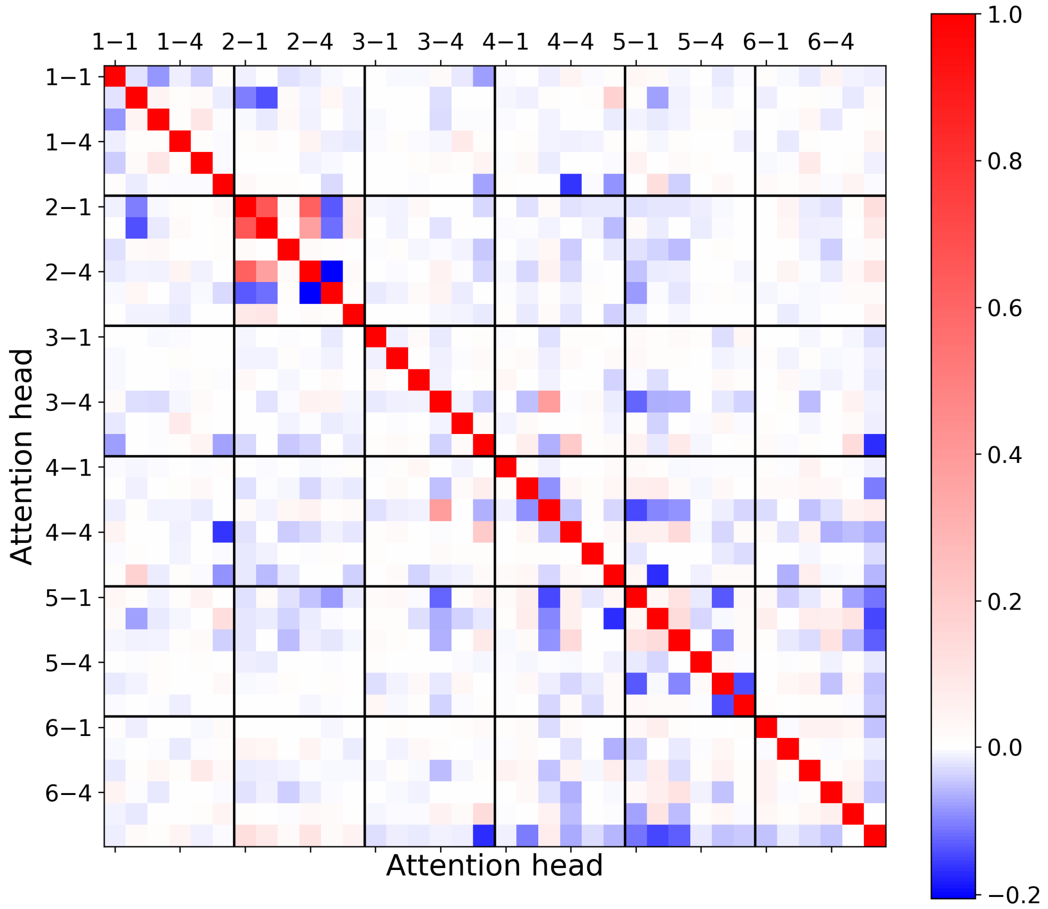


Figure S10: The correlation matrix between the weights attributed to the best scoring positions of each of the attention heads (exact positions given in Supplementary Figure S5 and S6). Attention heads are grouped by layer (first index).

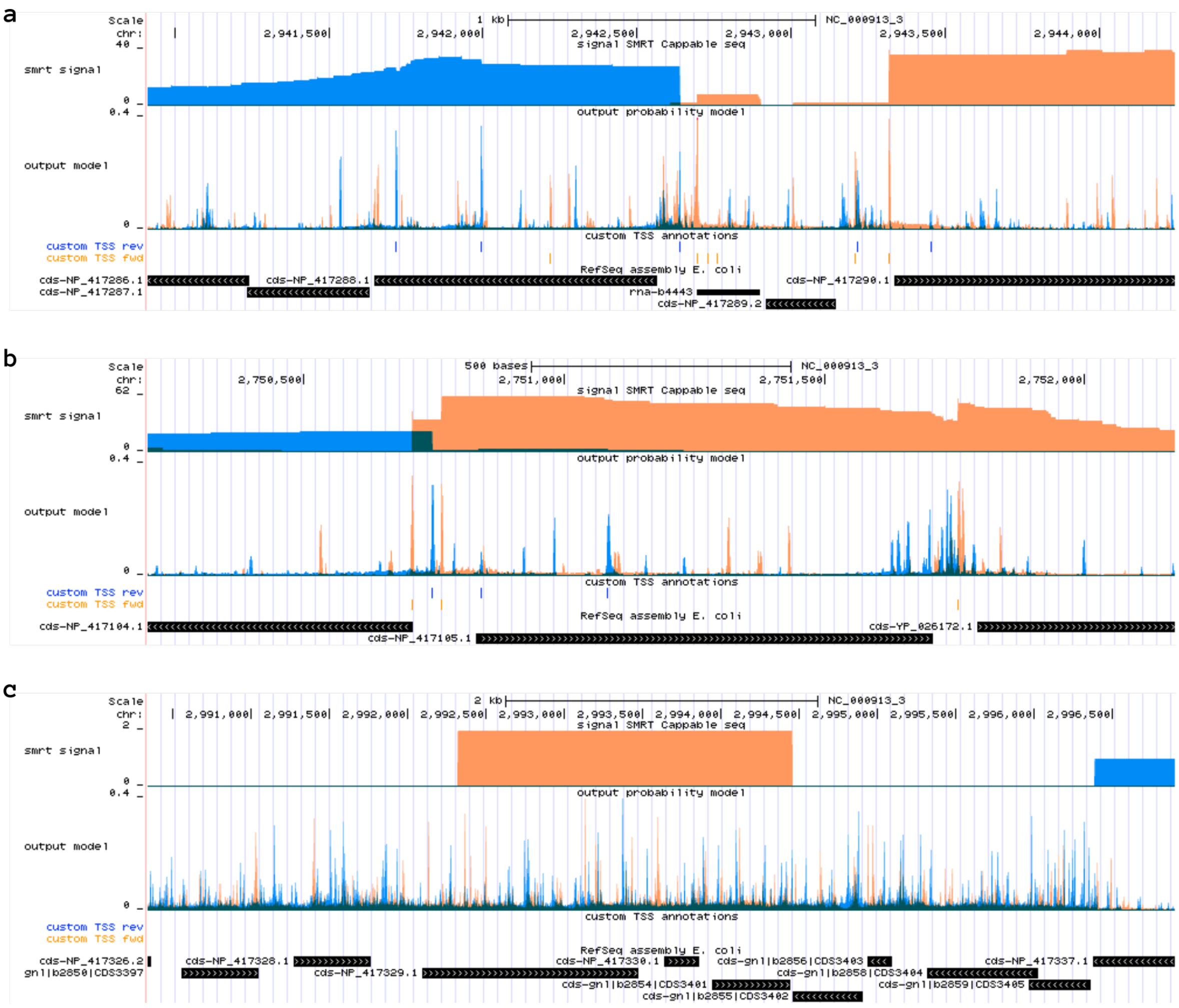


Figure S11: A view of the UCSC browser featuring the output predictions of the model, available at (`https://kermit.ugent.be/files/UCSC/UCSC_browser.html`). For each track is given, from top to bottom: the nucleotide position on the genome, the signal of the SMRT-Cappable-Seq experiment, the output of the model, the custom annotations of the test set for both the forward and reverse strand and the RefSeq assembly. Orange and blue are used to display both the SMRT-Cappable-seq signal and the model output probability for the forward and reverse strand, respectively. (**a**,**b**) give examples of increased activity of the antisense/sense strand for regions positioned between coding sequences. Regions furthermore highlight the consensus between the model activity and the signal of SMRT-Cappable-Seq. (**c**) Regions spanning pseudogenes are characterized by the continuous high output probability of the model on both sense and antisense.
